# RICO enables integrated analysis of ribosomal DNA copy number and methylation from nanopore long-read sequencing

**DOI:** 10.64898/2026.07.05.736662

**Authors:** Zaka Wing-Sze Yuen, Noemie Leeder, Thejaani Udumanne, Andrew Garvie, Lee Wong, Emiliana Weiss, Lex van Loon, Austen R.D. Ganley, Ross Hannan, Eduardo Eyras, Nadine Hein

## Abstract

Ribosomal RNA (rRNA) provides the structural and catalytic core of ribosomes and is encoded by ribosomal RNA genes (rDNA) arranged in tandem repeat arrays. rDNA copy number (CN) is highly dynamic, representing a clinically relevant form of structural variation, but its accurate quantification has been challenging due to its highly repetitive and GC-rich nature. Here, we present RICO (RIbosomal dna Integrated COpy number and methylation analysis), a novel computational pipeline for integrated estimation of rDNA CN and methylation using Oxford Nanopore long-read sequencing. RICO leverages long sequencing reads that span entire rDNA repeats, mapped to an rDNA-augmented reference genome, and normalizes coverage using a single-copy gene panel. We demonstrate that RICO provides accurate rDNA CN estimates in simulated datasets and reproducible measurements across human samples, with strong agreement to short-read sequencing and PCR-based methods. Furthermore, RICO reproduces a ∼40% reduction in rDNA CN in *Atrx*-knockout mouse embryonic stem cells, consistent with established effects of ATRX loss on rDNA CN. RICO also captures an increased total and active rDNA CN in malignant cells from a MYC-driven B-cell lymphoma mouse model, in agreement with prior studies. Applying RICO to independent human cohorts, we uncover that individuals with higher total rDNA CN consistently exhibit higher fractions of highly methylated rDNA copies, suggesting a dosage compensation mechanism that potentially maintains a similar number of active rDNA copies across individuals. Together, RICO enables integrated analysis of rDNA CN and methylation state, providing a scalable framework for investigating rDNA regulation across population and disease studies. RICO is available at https://github.com/comprna/RICO.

## Introduction

Ribosomal RNAs (rRNAs) are encoded by ribosomal RNA genes (rDNA) and form the structural and catalytic core of ribosomes, the essential molecular machines responsible for protein synthesis. In mammalian genomes, rDNA is organized as tandem repeat arrays distributed across multiple chromosomes (Fig. 1a). Each ∼43 – 45 kb repeat contains a single transcription unit encoding the 18S, 5.8S, and 28S rRNAs, which are transcribed as a 45S precursor rRNA by RNA polymerase I (Pol I), representing the first and rate-limiting step of ribosome biogenesis (Moss, Langlois, Gagnon-Kugler, & Stefanovsky, 2007; Panov, Hannan, Hannan, & Hein, 2021). Adjacent transcription units are separated by an approximately 30 kb intergenic spacer (IGS), which contains regulatory elements controlling rDNA transcription and repeat stability

**Figure 1.**
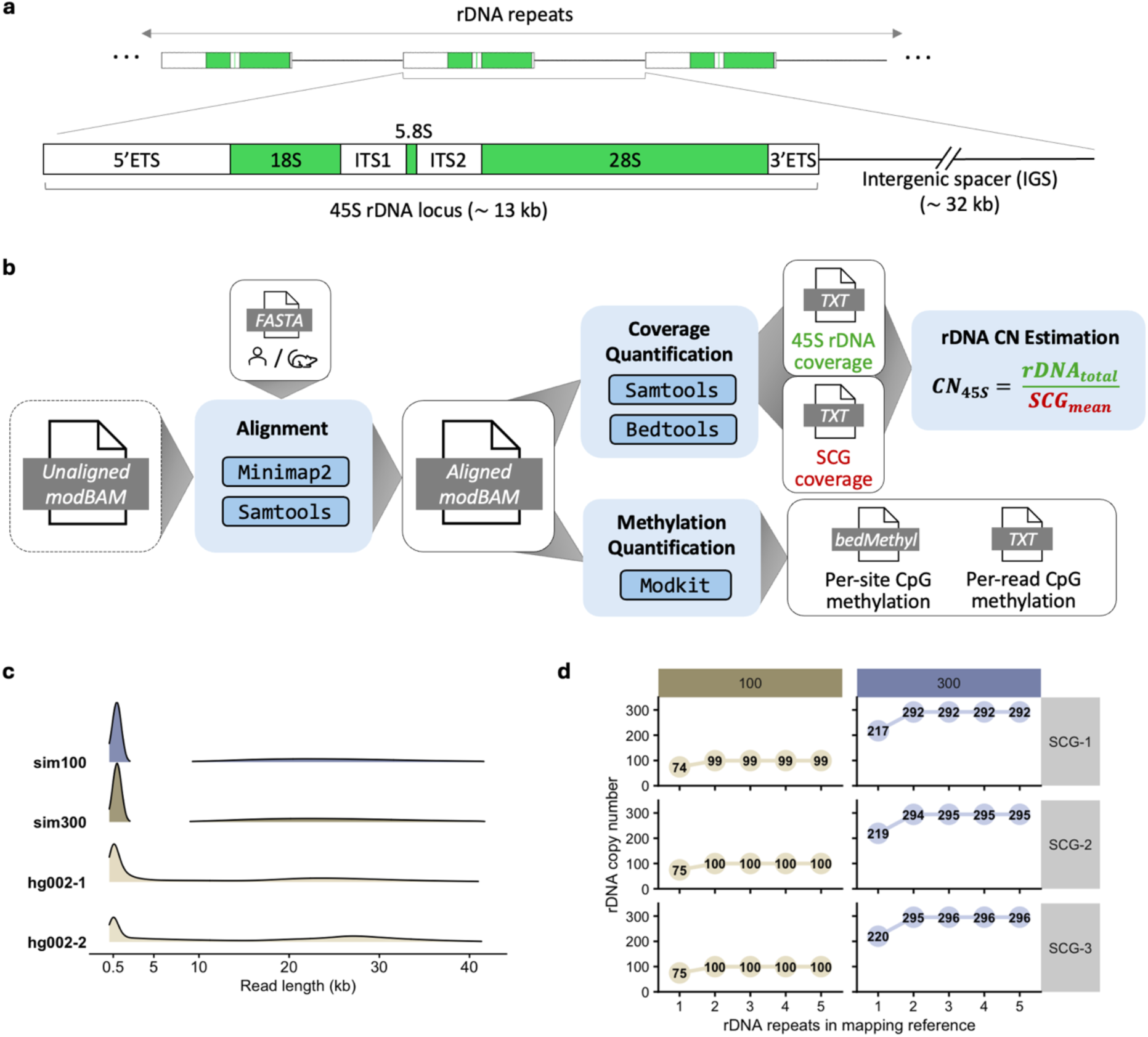
RICO accurately recovers predefined rDNA copy numbers (CN) in simulated long-read datasets. (**a)** Schematic representation of rDNA repeat organization in mammalian genomes. rDNA repeat units are arranged in long tandem arrays. Each repeat is ∼43-45 kb in length and contains a single 45S rDNA locus, comprising the 18S, 5.8S, and 28S rRNA coding regions (shown in green) separated by internal transcribed spacers (ITS1 and ITS2) and flanked by external transcribed spacers (5ʹETS and 3ʹETS), followed by an intergenic spacer (IGS). **(b)** Overview of RICO for estimating rDNA CN from long-read Nanopore sequencing data. RICO takes an unaligned modified BAM (modBAM) file as input, which can be generated by basecalling raw signal data, i.e. POD5, using Nanopore basecallers such as *Dorado*. Unaligned modBAM files are aligned to human or mouse reference genomes containing five concatenated rDNA repeats using *minimap2* and *samtools*. The human rDNA reference KY962518.1 and mouse rDNA reference BK000964.3 are used for human and mouse samples, respectively (see Methods). After read alignment, only primary alignments are retained to ensure reads are uniquely mapped to either the rDNA or non-rDNA regions of the genome, reducing contributions from rDNA-like regions elsewhere in the genome. This produces an aligned modBAM file that serves as the basis for both coverage quantification and methylation analysis. Coverage across the 45S rDNA locus and curated panels of single-copy genes (SCGs) is quantified using *samtools* and *bedtools*. For references containing multiple concatenated rDNA copies, mean coverage values from individual rDNA units are summed to obtain total rDNA coverage, and rDNA CN is estimated as the ratio of summed rDNA coverage to mean SCG coverage. In parallel, CpG methylation is quantified directly from aligned modBAM files using *modkit*, enabling joint analysis of rDNA CN and rDNA methylation state. RICO is implemented as both a Nextflow-based high-performance computing workflow and a standalone bash workflow for local execution and is available at https://github.com/comprna/RICO. **(c)** Read-length distributions of simulated datasets (sim100 and sim300) and two PromethION whole-genome sequencing runs of HG002 (hg002-1 and hg002-2). Simulations were parameterised to recapitulate the bimodal read length distribution observed in HG002 data, with these simulated short-read and long-read length profiles being combined. **(d)** Estimated rDNA CN across different number of rDNA repeats (1 to 5) included in the mapping reference for simulations with 100 (left) and 300 (right) known rDNA copies. CN were calculated using the mean-coverage approach with three different single-copy gene (SCG) reference panels (SCG-1, SCG-2, SCG-3). Results for the mode-coverage approach are shown in Supplementary 1.

In humans, rDNA repeats are located within the nucleolar organizer regions (NORs) on the short arms of the five acrocentric chromosomes (13, 14, 15, 21, and 22), whereas in mice they are located on chromosomes 12, 15, 16, 18, and 19 (Britton-Davidian, Cazaux, & Catalan, 2012; Henderson, Warburton, & Atwood, 1972; McStay, 2016). Despite their essential role in protein synthesis, the number of rDNA repeats, referred to as the rDNA copy number (CN), varies substantially between individuals, ranging from approximately 100 to 600 copies per haploid genome (Francisco Rodriguez-Algarra, Evans, & Rakyan, 2024). This variation reflects the dynamic nature of rDNA CN, although the mechanisms underlying rDNA CN variation in mammals remain unclear. Consequently, rDNA CN represents a highly variable form of structural variation whose biological significance remains incompletely understood.

Accumulating evidence suggests that rDNA CN is more dynamic than traditionally assumed and can change in response to genetic, developmental, or environmental factors. Altered rDNA CN has been associated with a wide range of biological conditions, including Bloom syndrome and other DNA repair disorders, cancer, and ageing-related phenotypes (Gál et al., 2024; Hallgren, Pietrzak, Rempala, Nelson, & Hetman, 2014; Hori, Engel, & Kobayashi, 2023; Valori et al., 2020). For instance, loss of Alpha-Thalassemia/X-linked Intellectual Disability *(ATRX)*, a chromatin-modifier and tumour-suppressor gene frequently mutated in gliomas and other malignancies, results in significant rDNA CN reduction and increases tumour cell sensitivity to RNA Pol l inhibitors (Udugama et al., 2018). However, despite these associations, it remains unclear whether changes in rDNA CN directly contribute to disease phenotypes or instead reflect broader alterations in genome maintenance and cellular homeostasis.

One explanation for inconsistent relationships between total rDNA CN and biological phenotype is that only a subset of repeats is transcriptionally active/competent at any given time. rDNA repeats exist in distinct chromatin states, with transcriptionally active repeats generally exhibiting low DNA methylation, whereas inactive repeats are typically DNA hypermethylated (Gagnon-Kugler, Langlois, Stefanovsky, Lessard, & Moss, 2009; McStay & Grummt, 2008). Although rDNA methylation is not a direct measure of rRNA synthesis, it is widely used as a proxy for the transcriptional competence of individual rDNA copies in particular methylation at the regulatory and transcribed region. Consequently, individuals with similar total rDNA CN may differ substantially in the number of transcriptionally active/competent rDNA copies.

Previous studies have proposed that epigenetic regulation may buffer variation in total rDNA CN by maintaining a relatively stable pool of transcriptionally active/competent rDNA repeats (Hori et al., 2023; Hori, Shimamoto, & Kobayashi, 2021; Parks et al., 2018; F. Rodriguez-Algarra et al., 2022). If true, this model would explain why substantial variation in total rDNA CN is often tolerated without obvious phenotypic consequences. However, this hypothesis has not been systematically evaluated across diverse mammalian samples as existing methods have limitations in simultaneously quantifying total rDNA CN and the epigenetic state of individual rDNA repeats from the same sample.

Several experimental and computational methods have been developed to estimate rDNA CN, including quantitative real-time PCR (qPCR), digital droplet PCR (ddPCR), pulse-field gel electrophoresis (PFGE) and computational analyses of whole-genome sequencing (WGS) data. Experimental approaches generally require dedicated assays and are difficult to scale to large cohorts, whereas computational methods can leverage existing sequencing datasets. Existing computational approaches rely almost exclusively on short-read sequencing, where uneven coverage (i.e. drop-outs) across the highly repetitive and GC-rich rDNA repeat, together with mapping ambiguity, can bias CN estimation. These limitations have contributed to inconsistent rDNA CN estimates between studies and continue to complicate interpretation of disease-associated variation.

To address these limitations, we developed RICO, a novel computational pipeline for integrated estimation of rDNA CN and DNA methylation from Oxford Nanopore long-read WGS data. By leveraging long reads that span entire rDNA repeats, RICO reduces coverage biases associated with short-read sequencing and improves repeat estimate accuracy, while simultaneously enabling methylation analysis within the same sample. This integration enables simultaneous estimation of both total rDNA CN and transcriptionally active/competent rDNA. We benchmark RICO using simulated datasets with predefined CN, independent sequencing runs of reference samples, and short-read-derived rDNA CN estimates from the 1000 Genome Project. We further validate RICO in dynamic biological systems using mouse models with established alterations in rDNA CN, including *Atrx*-knockout (KO) mouse embryonic stem cells (mESCs) with expected ∼40% rDNA CN reduction and MYC-driven lymphoma cells with increased transcriptional active rDNA repeats. Across human samples, integrated CN and methylation analyses with RICO revealed a strong inverse relationship between total rDNA CN and the fraction of transcriptionally active/competent rDNA repeats, suggesting a potential buffering mechanism that may help maintain a relatively stable pool of active rDNA copies despite variation in total CN. Together, our results demonstrate that RICO enables the integrated analysis of rDNA CN and methylation, providing a more complete view of rDNA regulation in normal and disease contexts.

## Results

### RICO accurately recovers pre-defined rDNA copy numbers from simulated long-read datasets

RICO is a computational framework that uses long-read Nanopore WGS data to quantify rDNA CN while retaining per-read CpG methylation information (Fig. 1b). RICO aligns Nanopore reads to species-specific reference genomes augmented with tandem rDNA repeat units and estimates rDNA CN by normalising coverage across the 45S rDNA locus to curated single-copy gene (SCG) reference panels. The same aligned reads are subsequently used for methylation profiling, enabling rDNA CN and methylation state to be characterised from a single sequencing dataset. Because Nanopore reads can span an entire repeat, and in some cases multiple consecutive repeats, RICO was designed to accommodate reads extending across individual repeat boundaries while reducing ambiguity arising from highly homologous rDNA-like sequences elsewhere in the genome.

To evaluate RICO, we first used simulated long-read Nanopore datasets with predefined rDNA CN to optimise the rDNA reference configuration and evaluate the accuracy of the coverage-based CN estimation strategy. We simulated Nanopore reads using *Badread (Wick, 2019)* from a human reference genome with a fixed number of rDNA repeats in a single repeat array, considering two possible configurations of 100 and 300 repeats. Simulation parameters were informed by publicly available Nanopore WGS data from the human reference sample HG002, which exhibited a bimodal read-length distribution that was recapitulated in the simulated benchmarking datasets (Fig. 1c) (Methods).

We used these simulated datasets to evaluate the following three components of the RICO CN estimation strategy: (1) the number of rDNA repeat units required in the mapping reference for accurate CN estimation, (2) the choice of SCG panel for coverage normalisation, and (3) the metric to summarise sequencing coverage across repeats. Augmented mapping references containing between one and ten concatenated rDNA repeat units in a single array were evaluated. Additionally, as normalisation controls, we compared three independently curated human SCG panels (SCG-1, SCG-2 and SCG-3), generated using increasingly stringent thresholds for sequence-uniqueness filtering and differing in the number of retained genes (please see Methods for further detail). We also compared a mean-coverage approach implemented in RICO with a previously described mode-based approach described by Sharma et al. (2022) for short-read sequencing data.

In both simulations, CN estimates increased with the number of rDNA repeats included in the mapping reference and plateaued at approximately the expected CN when more than two repeats were included (Fig. 1d), indicating that mapping saturation had been reached; hence, expanding the mapping reference with additional rDNA repeats would not affect coverage estimates. Secondly, we found that while CN estimates were broadly consistent across SCG panels, normalization to SCG-2 and SCG-3 yielded estimates closest to the predefined CN (Fig. 1d). Thirdly, the mean-coverage method consistently produced estimates closer to the expected values than the mode-coverage approach (Fig. S1), suggesting that mean coverage better captures quantitative differences in rDNA abundance. Overall, the simulations demonstrate that RICO can reliably recover predefined rDNA CN under controlled conditions and support the use of mean-coverage for subsequent CN estimation.

### RICO recovers consistent rDNA copy number estimates across independent sequencing experiments

We next evaluated the reproducibility of RICO using publicly available Nanopore WGS data from the Genome in a Bottle (GIAB) Ashkenazi Trio (HG002, HG003 and HG004) and Han Chinese trio (HG005, HG006 and HG007). Consistent with the simulated results, rDNA CN estimates for the Ashkenazi Trio stabilised when two or more rDNA repeats were included in the augmented mapping reference (Fig. 2a; Fig. S2-S3). CN estimates were highly concordant between independent sequencing runs, with HG002 yielding 231 and 236 copies, HG003 yielding 182 and 184, and HG004 yielding 243 and 244 (Fig. 2b).

**Figure 2.**
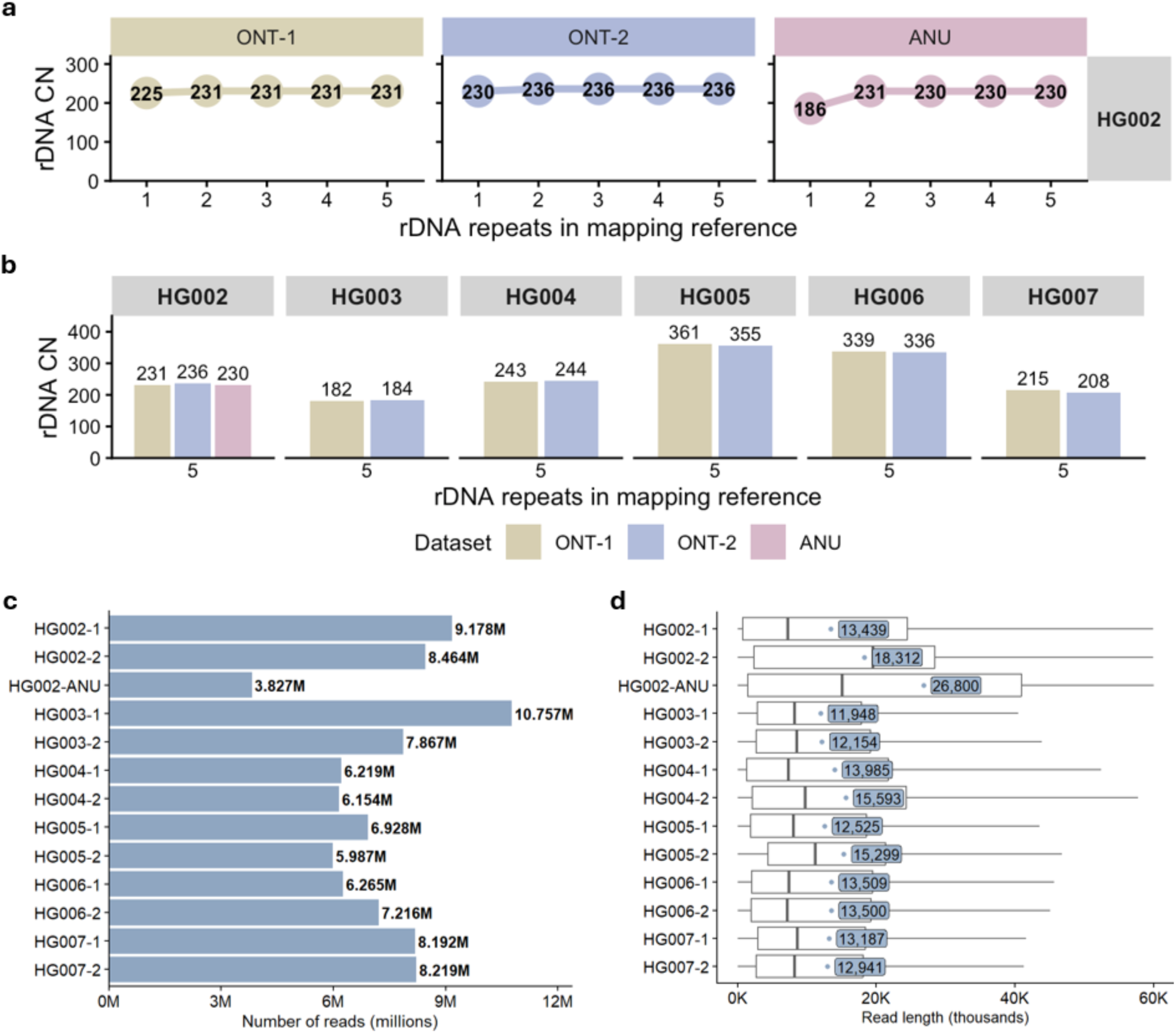
RICO produces reproducible rDNA CN estimates across independent sequencing datasets. **(a)** Estimated rDNA CN (y-axis) as a function of the number of rDNA repeats included in the mapping reference (x-axis; 1-5 repeats) for the Ashkenazi trio samples HG002 (son). Results are shown for two independent Oxford Nanopore sequencing datasets (ONT-1: left and ONT-2: right) with the in-house sequencing sample (HG002-ANU: right). CN estimates were generated using the mean-coverage approach and the single-copy gene panel 2 (SCG-2). **(b)** rDNA CN estimates for the GIAB Ashkenazi trio (HG002, HG003 and HG004) and Han Chinese trio (HG005, HG006 and HG007) derived from two independent Oxford Nanopore sequencing datasets (ONT-1 and ONT-2). An additional independently generated Nanopore WGS dataset for HG002 (ANU) was included to further assess reproducibility across sequencing experiments. CN estimates were generated using the mean-coverage approach with a mapping reference containing five concatenated rDNA repeats and single-copy gene panel 2 (SCG-2). Benchmarking across different numbers of rDNA repeats in the mapping reference, SCG panels, and mean- and mode-coverage approaches is shown in Supplementary Figs. S2-S4. **(c)** Total number of reads generated for each dataset of the trio samples, obtained from the unaligned modified BAM (modBAM) files. Values shown to the right of each bar indicate the total number of reads (in millions) per dataset. **(d)** Read-length distributions for each sequencing dataset of the trio samples, derived from unaligned modBAM files. Boxplots summarise the read length distribution, with blue dots and accompanying labels indicate the mean read length for each dataset.

To further assess reproducibility across independently generated sequencing data, we performed an additional Nanopore WGS experiment from HG002 cells (HG002-ANU). RICO estimated 230 rDNA copies, closely matching estimates obtained from the publicly available HG002 sequencing runs (Fig. 2a-b; Fig. S4). The rDNA CN estimates remained highly consistent across datasets, despite differences in sequencing yield and read-length distributions, including a lower read yield (∼3.8M reads) and longer mean read length (∼26 kb) in HG002-ANU compared with the publicly available datasets (∼6-10M reads and ∼13 kb mean read length) (Fig. 2c-d). Together, these results demonstrate that RICO yields reproducible rDNA CN estimates across independently generated Nanopore sequencing datasets generated from the same cell line. Analysis using the alternative mode-coverage approach yielded generally similar CN estimates but showed increased run-to-run and SCG-dependent variability relative to the mean-coverage method (Fig. S3-S4), further supporting the use of the mean-coverage approach.

Given the stable CN estimates obtained with two or more concatenated rDNA repeats, we used an augmented reference containing five repeat units for all subsequent analyses. Applying RICO to the Han Chinese trio similarly produced highly concordant CN estimates between independent sequencing runs, with differences of no more than seven copies for each sample (Fig. 2b). Across both trios, differences between replicate CN estimates ranged from one to seven copies, further demonstrating the reproducibility of RICO across independently generated sequencing datasets.

### RICO accurately recovers established biological and population-level variation in rDNA copy number

To test whether RICO can detect biologically established rDNA CN differences, we studied a mESC model with experimentally validated rDNA CN loss. Knockout (KO) of the *Atrx* gene, a tumour suppressor frequently mutated in gliomas and other cancers and linked to genome instability, was previously shown to result in a relative 40-50% reduction in rDNA copies in the *Atrx*-KO compared to wild-type (WT) mESCs (Udugama et al., 2018).

We performed Nanopore WGS on WT and *Atrx*-KO mESCs and applied RICO to estimate rDNA CN. As RICO was initially developed and optimised using human datasets, we first assessed whether CN estimates were similarly stable across mouse augmented references containing between one and ten concatenated rDNA repeats (Fig. S5). Mean-based CN estimates remained consistent across all reference configurations (Fig. S5). Importantly, we identified a 39% reduction in rDNA CN in *Atrx-*KO relative to WT mESCs across all configurations (122 vs. 74 copies) (Fig. 3a; Fig. S5). These DNA CN closely match the previously reported relative estimates determined by qPCR, highlighting the accuracy of the RICO pipeline.

**Figure 3.**
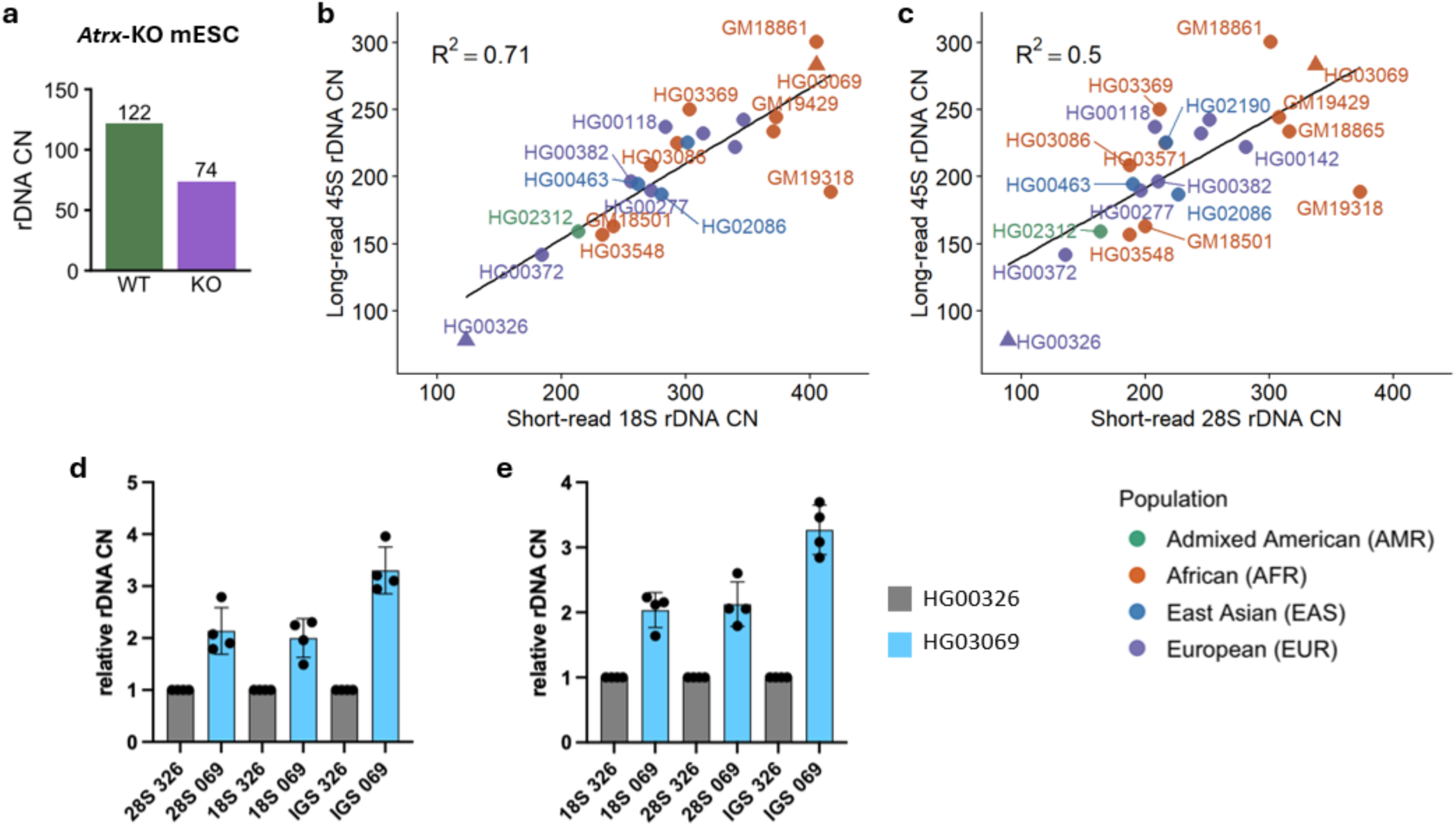
RICO detects biologically established changes in rDNA copy number (CN). **(a)** Estimated rDNA CN for wild-type (WT) and *Atrx*-KO mouse embryonic stem cells (mESCs). Benchmarking across different numbers of rDNA repeats in the mapping reference and mean- and mode-coverage approaches is shown in Supplementary Fig. S6. Read-length distributions for these samples are shown in Supplementary Fig. S7. **(b-c)** Comparison of RICO long-read rDNA CN estimates to short-read-based rDNA 18S **(a)** and 28S **(b)** CN estimates across individuals. Black lines show linear regression fit, with the corresponding r-squared (R^2^) values shown. HG00326 and HG03069, highlighted by triangles, were selected for subsequent qPCR validation. **(d-e)** qPCR validation of rDNA CN differences between HG00326 and HG03069. Relative abundance of the 18S, 28S, and IGS regions was quantified by qPCR, normalised to either **(d)** *RPPH1* or **(e)** *TP53*. Grey bars represent HG00326 and light-blue bars represent HG03069. Points indicate replicates from the same cell line isolated at different passage number (n = 4), and bars represent indicate ± SD.

To further validate the applicability of RICO for consistent rDNA CN estimations across independent datasets and sequencing platforms, we next examined samples from the 1000 Genomes Project (1KGP). We compared RICO estimates derived from Nanopore long-read data with previously published short-read-derived rDNA CN estimates from Hall, Turner, and Queitsch (2021) to assess cross-platform reproducibility across 1KGP samples. Although short-read-based estimates have limitations, they remain widely used for population-level analyses, as they provide consistent relative differences in rRNA genes between individuals (Hall et al., 2021) and thus provide a reference point for validating the consistency of RICO rDNA CN estimates. In total, we analyzed 22 individuals from four different ancestry groups (African, American, East Asian, and European) for which we identified a set of short-read (Hall et al., 2021) and Nanopore long-read data (Gustafson et al., 2024).

To assess concordance, we estimated 45S rDNA CN from the Nanopore data using RICO and compared these estimates with the available short-read CN estimates for the 18S and 28S regions, which were derived by normalising rDNA read depth to chromosome 1 coverage (Hall et al., 2021). We found that long- and short-read estimates were positively correlated, with short-read estimates being generally higher than long-reads ones (Fig. S6). Short-read 18S CN explained 71% of the variance in RICO-derived 45S CN (Fig. 3b), while short-read 28S CN explained 50% of the variance (Fig. 3c).

To further validate RICO estimates, we experimentally tested rDNA CN using qPCR in two of the cell lines, HG00326 and HG03069, for which there were markedly different rDNA CN estimates confirmed by both long-read and short-read approaches (Fig. 3b-c). Consistent with both sequencing-based approaches, HG03069 showed substantially higher relative CN across 18S, 5.8S, 28S, and IGS regions than HG00326 (Fig. 3d-e), a result that was robust to normalization with two different single-copy genes, *RPPH1* and *TP53*. Thus, our RICO long-read analysis recapitulates a significant proportion of the relative differences in rDNA CN between individuals that are detected by short-read analysis, and can be validated by qPCR. Together, these results indicate that RICO yields stable and biologically consistent rDNA CN estimates under conditions with experimentally validated rDNA CN alterations.

### rDNA methylation profiles are reproducible yet highly variable between individuals

Having established that RICO provides robust estimates of total rDNA CN, we next asked how rDNA CN relates to the methylation state of individual rDNA repeats. Although rDNA CN varies substantially between individuals, the extent to which this variation translates into differences in the number of transcriptionally active/competent rDNA repeats remains unclear. As rDNA methylation is associated with transcriptional state, integrating methylation profiling with CN estimation provides a means to distinguish variation in total rDNA abundance from variation in the putatively transcriptionally active/competent repeat fraction.

We first examined whether methylation profiles across the transcribed regions of rDNA were highly reproducible across sequencing runs. To classify methylation at the individual-repeat level, CpG calls and rDNA reads from transcribed regions were filtered based on calling confidence and number of CpG sites per read, respectively, and the proportion of 5mC calls was calculated for each retained read (Fig. 4a; Methods).

**Figure 4.**
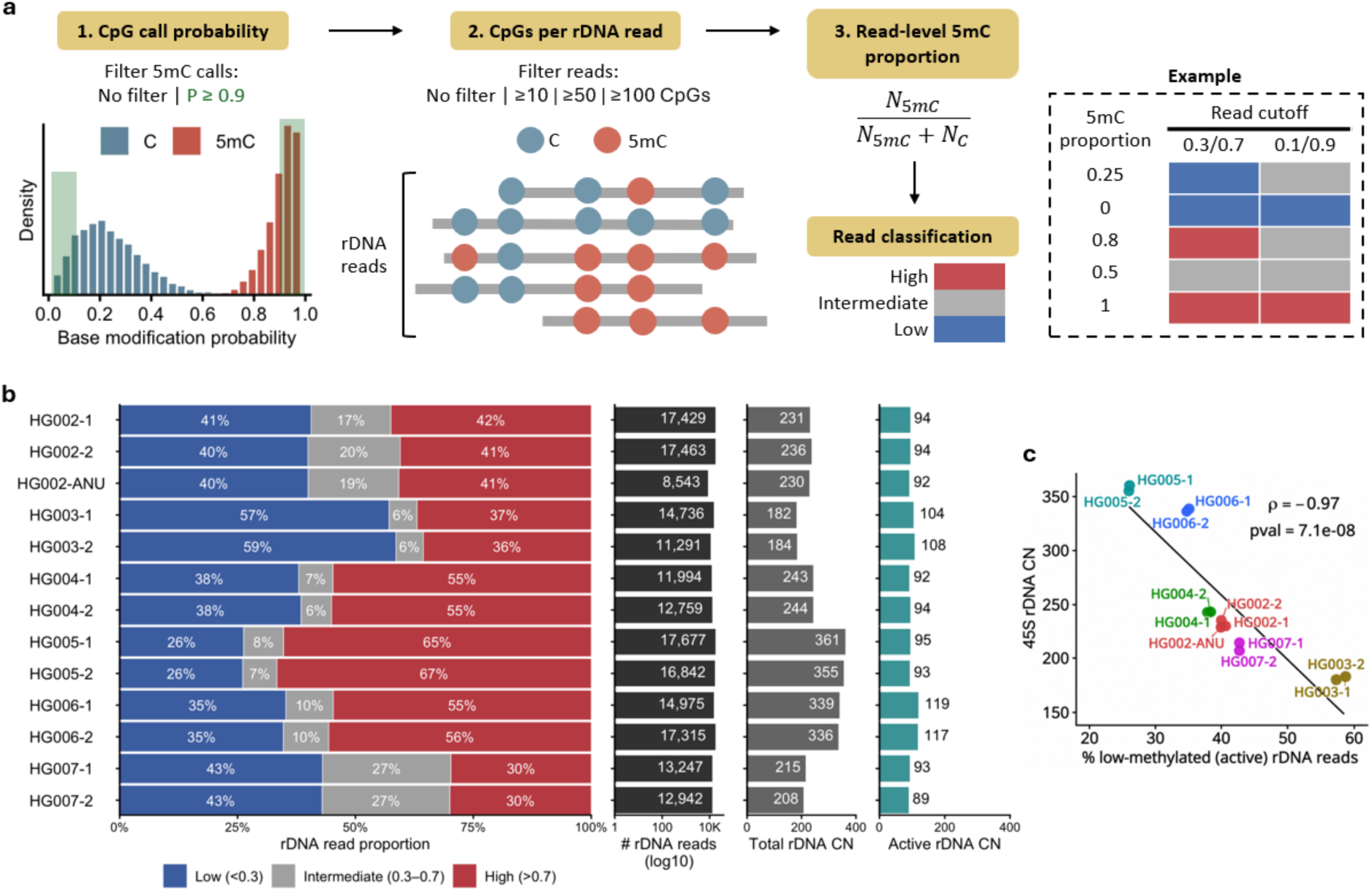
rDNA methylation profiles are reproducible and reveal constrained active rDNA repeat abundance: The GIAB trio methylation and the active CN estimates. **(a)** Schematic of the read-level rDNA methylation classification workflow. The left barplot shows the distribution of probability per site in individual reads provided by dorado, separated by methylated (red) and non-methylated (blue). Within each individual rDNA read, we follow two strategies, either considering all predicted CpG sites (all calls) or only those with probability ≥0.9. We also considered different strategies depending on the number of CpG sites retained per read, considering all reads regardless of their number of CpGs (all), or only reads with ≥10, ≥50, or ≥100 CpGs. In each configuration, we then calculate the 5mC proportion as the number of 5mC calls divided by the total number of 5mC and canonical C calls. Reads were subsequently classified as low, intermediate, or highly methylated using methylation-proportion cutoffs of either 0.3/0.7 or 0.1/0.9. The example illustrates how individual reads can be assigned differently depending on the classification cutoffs applied. **(b)** Per-read rDNA methylation profiles across samples using 0.3/0.7 classification cutoffs. Each horizontal bar represents the proportion of rDNA reads classified as low (<0.3), intermediate (0.3-0.7), or high (>0.7) methylation based on the proportion of 5mC calls per read. All CpG calls were included regardless of calling probability (i.e. no filtered), and no minimum number of CpG sites per read (i.e. no filtered) was required for read-level methylation classification. The accompanying panels show the total number of rDNA reads (log10 scale), estimated total rDNA CN, and estimated active rDNA CN calculated as the proportion of low-methylation reads multiplied by total rDNA CN. **(c)** Relationship between total rDNA CN (45S CN) and the proportion of low-methylation (active) rDNA reads (as determined from the proportion in blue in b) across samples. Each point represents an individual run. Colour represents the same sample with multiple sequencing runs. The black line shows the linear regression fit, with Spearman correlation (ρ) and corresponding p-value indicated.

Base modification calls showed distinct probability distributions for canonical C and 5mC across the trio samples (Fig. S8a). Applying a calling probability threshold of P ≥ 0.9 reduced the number of CpG calls, with a greater reduction in canonical C than in 5mC calls (Fig. S8b). We therefore examined the robustness of read-level methylation classification based on calling probability and read classification cutoffs. Per-read 5mC distributions remained broadly similar using either unfiltered 5mC calls or only high-confidence 5mC calls (P ≥ 0.9), with a greater proportion of reads falling between the classification cutoffs when unfiltered calls were used (Fig. S8c). Classification cutoffs of 0.3/0.7 generally fell within lower-density regions separating the low and high methylation populations, whereas the more stringent 0.1/0.9 cutoffs intersected high-density regions of the distributions in several samples (Fig. S8c). We therefore used 0.3/0.7 as the primary read-level classification thresholds for subsequent analyses.

Reads were then classified into three methylation states based on their 5mC proportion: low (<0.3), intermediate (0.3-0.7), and high (>0.7) methylation. The identification of an intermediate methylation population suggests that rDNA methylation states are not strictly binary. Interestingly, the proportion of all three methylation states was highly consistent across replicates from the same sample but varied substantially between individuals. For example, more than half of rDNA reads in HG003 were classified as low-methylation, whereas HG005 exhibited a substantially greater proportion of highly methylated reads (Fig. 4b). These results confirm the reproducibility of per-read rDNA methylation profiles across independent sequencing runs, and highlight individual-specific variation in rDNA methylation state.

To further examine the spatial distribution of methylation sites underlying these read-level classifications, we spatially mapped the individual C and 5mC calls on individual reads along a single 45S rDNA reference (Fig. S9). Low-methylation repeats were predominantly composed of canonical C calls across the 45S region, whereas highly methylated repeats showed widespread 5mC across the repeat (Fig. S9). HG007, which had the largest fraction of intermediate-methylated reads among the trio samples, exhibited a heterogeneous mixture of C and 5mC calls across the 45S region for the intermediate methylation state. While canonical C calls were more apparent around the 18S region, the reads do not present any spatial conservation of C or 5mC (Fig. S9). These patterns were highly reproducible between independent sequencing runs from the same individual, further supporting the reproducibility of the read-level methylation profiles.

Because low rDNA methylation is associated with transcriptionally permissive chromatin, we used the low-methylation fraction as a proxy for transcriptionally active/competent rDNA repeats. Intermediate methylated reads were not included in this estimate because their transcriptional state cannot be confidently assigned from methylation alone. We therefore estimated the number of putatively active rDNA copies by multiplying the total rDNA CN by the fraction of low-methylation reads for each sample (Fig. 4b, right panel). After averaging replicate sequencing runs for each individual, the estimated putatively active rDNA CN ranged from 90.5 to 118 copies, compared with 183 to 363 total rDNA copies (Fig. 4b; Table S1).

Across the trio samples, the total rDNA CN was strongly inversely correlated with the proportion of low-methylation rDNA reads (Spearman’s ρ = -0.97; Fig. 4c), such that individuals with higher total rDNA CN tended to have a smaller fraction of low-methylation repeats. This inverse relationship between total rDNA CN and the proportion of low-methylation reads was robust to variation in methylation classification parameters, including CpG calling probability thresholds, minimum numbers of CpG sites per read and alternative methylation cutoffs (Fig. S10-11). Methylation distributions remained consistent within samples across all parameter combinations (Fig. S10-11), and the inverse relationship between total rDNA CN and the proportion of low-methylation reads was maintained (Spearman’s ρ < -0.85) (Fig. S10-11). This inverse relationship suggests that epigenetic regulation buffers the variation in total rDNA CN by modulating the methylation of the 45S, thereby helping to maintain a relatively stable number of transcriptionally active/competent rDNA copies across individuals.

### Long-read methylation profiling captures known changes in transcriptionally active/competent rDNA repeats

We studied the ATRX model above to examine the rDNA methylation profiles at the single-read level, using methylation thresholds of <0.3, 0.3-0.7 and >0.7 to classify reads into low-, intermediate- and high-methylation states, respectively. Consistent with prior studies showing that rDNA in mESCs is predominantly unmethylated (Betto et al., 2021; Schlesinger, Selig, Bergman, & Cedar, 2009), all rDNA reads in both WT and *Atrx*-KO ESCs were classified to be in the low-methylation state (Fig. 5a).

**Figure 5.**
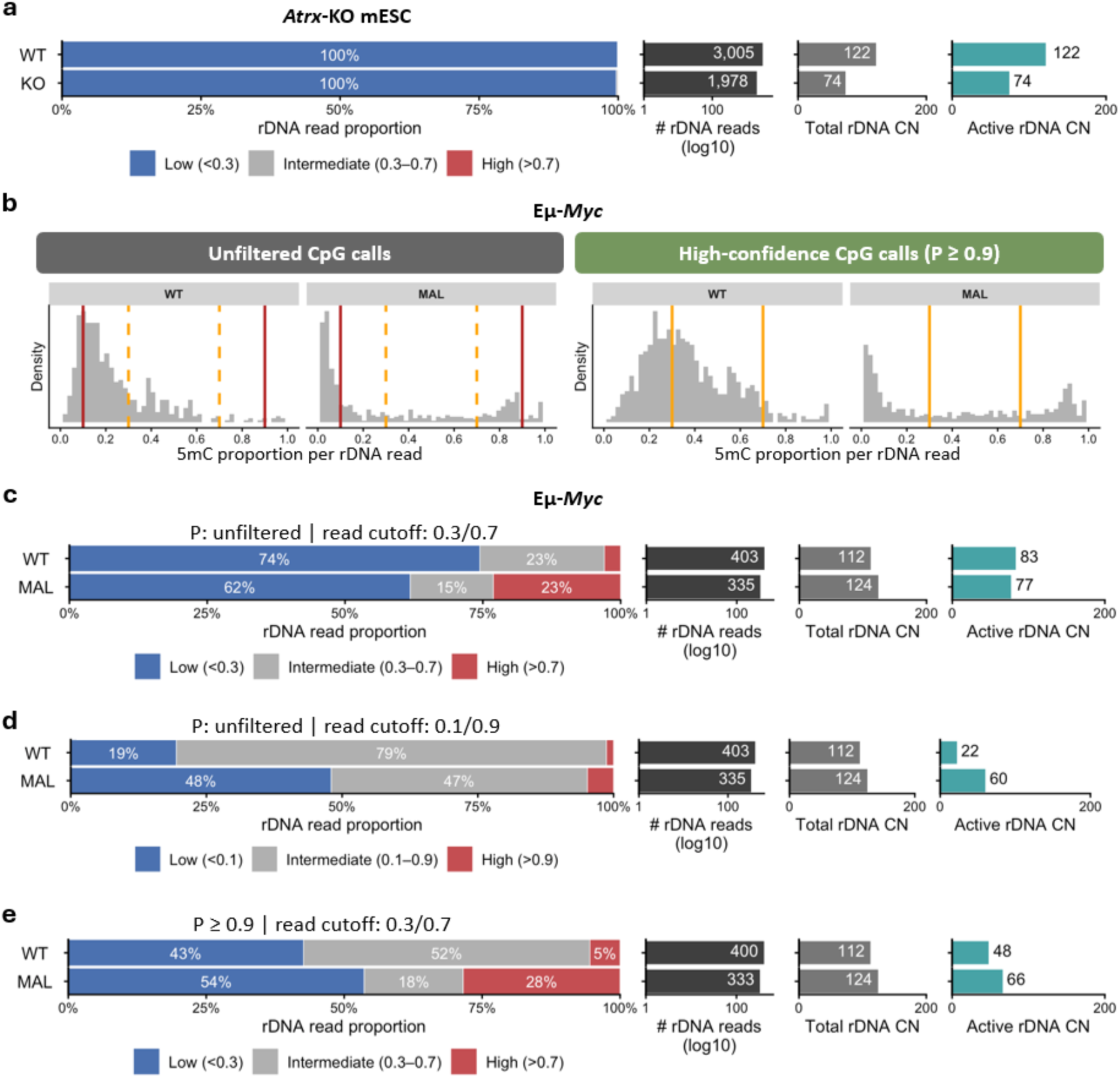
Methylation profiling recapitulates known active rDNA states in mouse models: ATRX methylation, *Eμ-Myc* methylation, active/competent copy number (CN) estimates (methylation validation). (a) Per-read rDNA methylation profiles in WT and *Atrx*-KO mESCs. Nearly all rDNA reads were classified as low-methylated (blue) in both WT and *Atrx*-KO cells. **(b)** Distribution of per-read 5mC proportions in WT and MAL Eμ-*Myc* cells using unfiltered CpG calls (left) or high-confidence CpG calls (P ≥ 0.9; right), with no minimum number of CpG calls required per read. Orange lines indicate the 0.3/0.7 classification cutoffs established in Fig. 4, while red lines indicate the alternative 0.1/0.9 cutoffs used to identify more extreme methylation states. **(c-e)** Per-read rDNA methylation profiles in WT and MAL Eμ-*Myc* cells using either (c) unfiltered CpG calls and 0.3/0.7 read-level classification cutoffs, (d) unfiltered CpG calls and 0.1/0.9 read-level classification cutoffs or (e) high-confidence CpG calls (P ≥ 0.9) and 0.3/0.7 read-level classification cutoffs. The total number of rDNA reads (log10 scale), total rDNA CN, and estimated putatively active rDNA CN (total rDNA CN × fraction of low-methylation reads) are shown in the adjacent panels.

Base-modification calling probabilities showed distinct distributions for canonical C and 5mC calls, with relatively few 5mC calls meeting the high-confidence probability threshold (P ≥ 0.9) (Fig. S12a & b). Despite this reduction, per-read 5mC proportion remained strongly concentrated within the low-methylation range under both unfiltered and high-confidence CpG calling conditions (Fig. S12c). Consistently, varying the minimum number of CpG calls required per read (≥10, ≥50 or ≥100), with or without the P ≥ 0.9 probability filter, had little effect on read classification. Consequently, almost all rDNA reads remained in the low-methylation state in both WT and *Atrx*-KO cells (Fig. S13). Consistent with this, spatial mapping of C and 5mC calls across individual rDNA repeats showed predominantly canonical C throughout the 45S region under both calling conditions (Fig. S14). These results indicate that the predominantly low-methylation state of mESC rDNA is robust to the filtering parameters used for read-level methylation detection.

Because the low-methylation fraction approached 100% in both samples, estimates of putatively active rDNA CN closely reflected their respective total CN estimates, yielding 122 copies in WT and 74 copies in *Atrx*-KO cells (Fig. 5a). These estimates were also largely insensitive to the alternative methylation filtering parameters, remaining close to the corresponding total rDNA CN estimates (Fig. S13). Therefore, the reduction in putatively active rDNA CN in *Atrx*-KO cells was driven primarily by the reduction in total rDNA CN rather than by a shift in the proportion of low-methylation repeats. This finding is consistent with prior results showing that *Atrx* loss decreases the abundance of transcriptionally active/competent rDNA chromatin (Udugama et al., 2018).

To provide further biological validation in a model with heterogeneous rDNA methylation, we applied RICO to Nanopore WGS data generated from a MYC-driven B-cell lymphoma mouse model (Eμ-*Myc*), in which malignant transformation is associated with elevated ribosome biogenesis, increased rRNA expression, and a significant increase in the fraction of transcriptionally active rDNA repeats in malignant (MAL) B-cells relative to WT controls, as previously demonstrated by psoralen cross-linking experiments (Bywater et al., 2012; Diesch et al., 2019). In contrast to mESCs, WT and MAL Eμ-*Myc* cells showed broad and multimodal distributions of per-read 5mC proportions (Fig. 5b), indicating substantially greater heterogeneity in rDNA methylation state.

Importantly, the inferred WT and MAL difference depended on how the low-methylation population was defined. Using unfiltered CpG calls and the 0.3/0.7 thresholds established in the human analysis, 74% of WT and 62% of MAL rDNA reads were classified as low-methylated (Fig. 5c). However, inspection of the continuous distributions showed that the MAL low-methylation population was more strongly concentrated near zero, whereas the WT distribution extended more broadly across the <0.3 range (Fig. 5b). We therefore examined a more stringent near-unmethylated category using a <0.1 threshold. Under this definition, 19% of WT compared with 48% of MAL reads were classified as low-methylated (Fig. 5d). Spatial mapping of individual methylation calls further showed that these near-unmethylated MAL repeats were dominated by canonical C calls across much of the 45S rDNA region (Fig. S16).

Modification call filtering also affected the methylation profiles in Eμ-*Myc* cells. Applying the high-confidence P ≥ 0.9 filter while retaining the 0.3/0.7 read thresholds shifted the per-read methylation distributions, particularly in WT cells (Fig. 5b), and resulted in 43% of WT compared with 54% of MAL reads being classified as low-methylated (Fig. 5e). This variability is consistent with the unequal retention of canonical C and 5mC calls after probability filtering (Fig. S15a-b), which alters the set of methylation calls contributing to each read-level 5mC proportion. Nevertheless, varying the minimum number of tested CpGs per read had very limited effects within each calling probability condition (Fig. S15c).

Together, these results demonstrate that, for samples with heterogeneous and multimodal rDNA methylation like the Eμ-*Myc* lymphoma cells, the read-level methylation classification cutoffs can obscure differences within the low-methylation population. Both the near-unmethylated fraction identified using unfiltered calls (i.e. P: unfiltered; <0.1) and the low-methylation fraction identified after high-confidence call filtering (i.e. P ≥ 0.9; <0.3) were increased in MAL relative to WT cells, concordant with the previously reported increase in transcriptionally active rDNA repeats in Eμ-*Myc* malignant B cells (Diesch et al., 2019). While methylation-based classification provides only a proxy rather than a direct measurement of transcriptional activity, the Eμ-*Myc* analysis demonstrates that the underlying distribution of read-level methylation states can reveal biologically relevant differences that are not captured by a single classification cutoff.

### Active/competent rDNA abundance is constrained across human populations

Our findings in the human trios prompted us to investigate whether the inverse relationships between total rDNA CN and proportion of low-methylated repeats were observed across a broader human cohort. We therefore examined rDNA methylation profiles across 1KGP samples (Fig. 6a).

**Figure 6.**
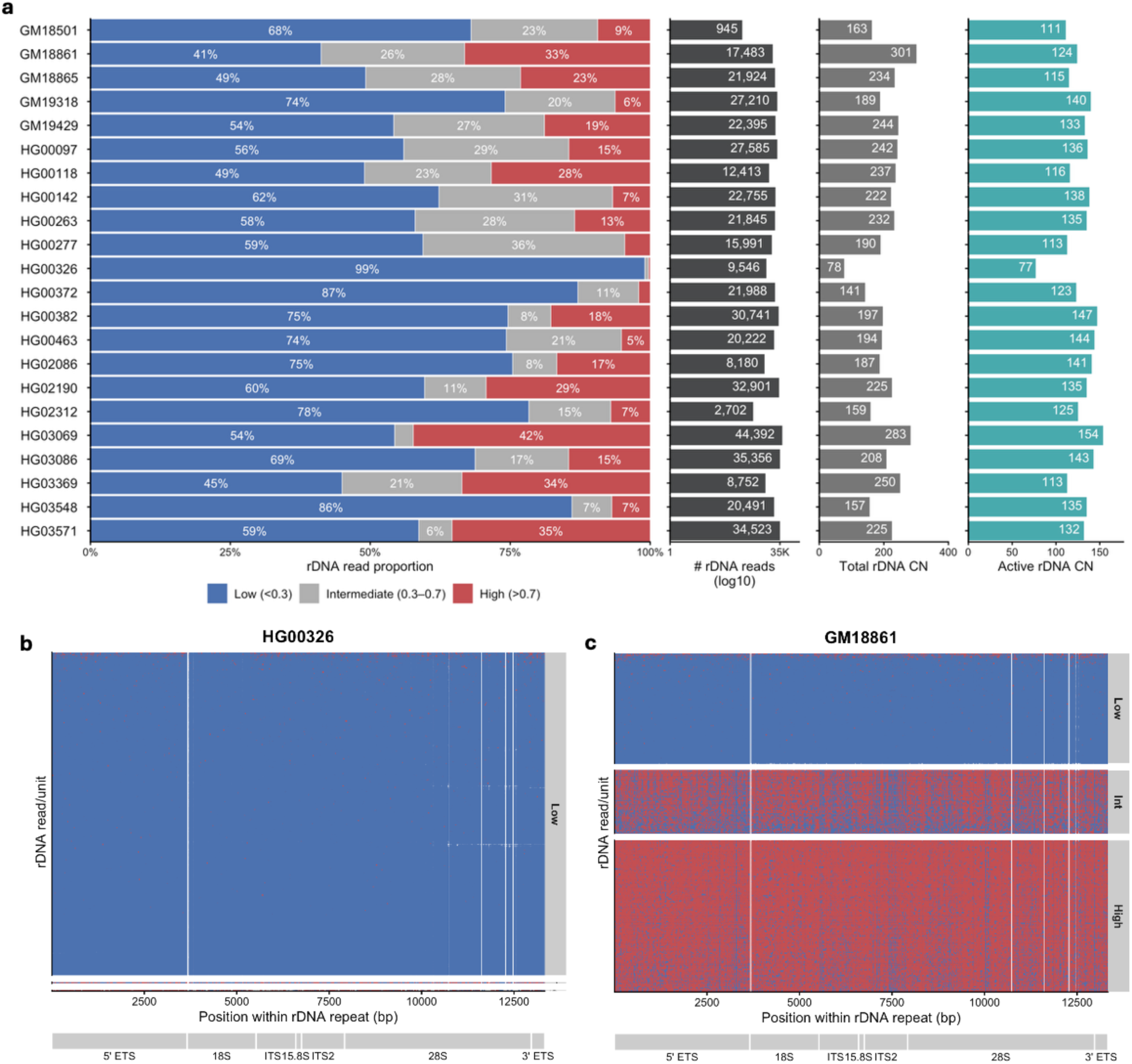
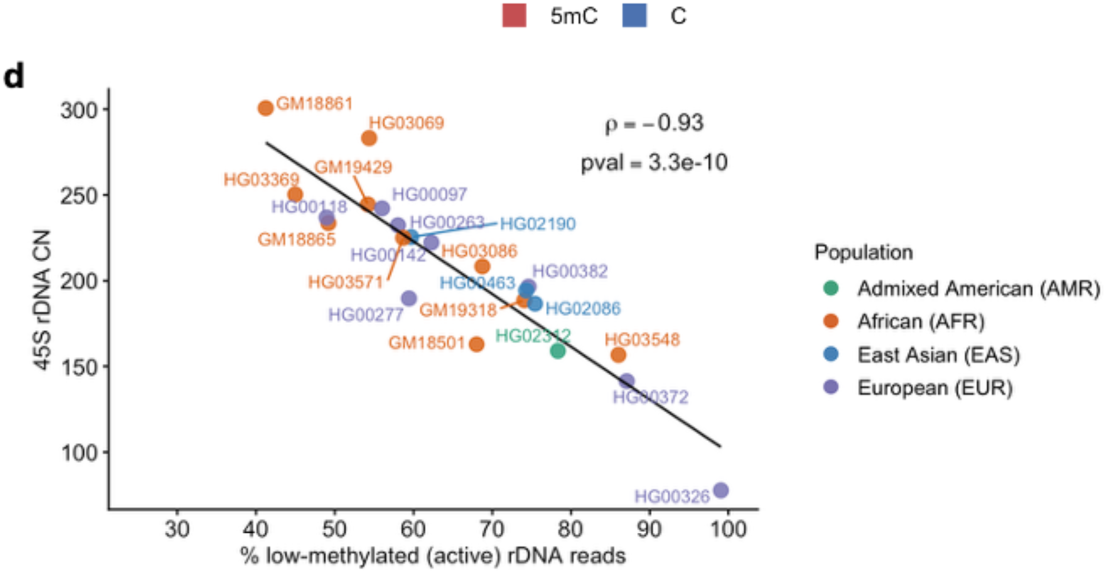
Epigenetic compensation constrains active rDNA copy number (CN) across 1KGP individuals with varying rDNA CN. **(a)** Per-read rDNA methylation profiles across samples. Each horizontal bar represents the proportion of rDNA reads classified as low (<0.3), intermediate (0.3 - 0.7), or high (>0.7) methylation based on the proportion of 5mC calls per read. Analyses were performed using all CpG calls and without filtering on the number of CpG sites per read. The total number of rDNA reads (log10 scale), total rDNA CN, and the estimated number of active rDNA copies (total rDNA CN x fraction of low-methylated reads) are shown in the adjacent panels. **(b-c)** Spatial distribution of canonical C and 5mC calls across individual 45S rDNA repeats in (b) HG00326 and (c) GM18861. Canonical C (blue) and 5mC (red) calls are shown according to their positions within the 45S rDNA repeat. Read/repeat units are grouped according to their overall 5mC proportion as low, intermediate, or high methylation. Spatial methylation profiles for all 1KGP samples are shown in Supplementary Fig. S17. **(d)** Relationship between total rDNA CN and the proportion of low-methylated (active) rDNA reads (from a, left panel) across samples. Each point represents an individual and is coloured by population affiliation. The black line shows the linear regression fit, with Spearman correlation (ρ) and corresponding p-value indicated.

Per-read rDNA methylation distributions varied substantially between individuals (Fig. 6a). Some samples, such as HG00326 and HG00372, exhibited predominantly low-methylation rDNA reads (99% and 87%, respectively), whereas others, including GM18861 and HG03069, showed substantially larger fractions of high-methylation reads (33% and 42%, respectively). Examination of individual C and 5mC calls across the 45S rDNA repeat further illustrated these differences. Low-methylation repeats were predominantly composed of canonical C calls across the 45S locus (Fig. 6b), whereas high-methylation reads showed widespread 5mC, with intermediate-methylated repeats displaying heterogenous mixtures of the two methylation patterns (Fig. 6b) (Fig. S17). These spatial profiles confirmed that the read-level methylation classification reflected distinct patterns of C and 5mC calls across individual 45S rDNA repeats (Fig. 6b-c; Fig. S17).

We then considered the low-methylation reads, as before, to estimate transcriptionally active/competent rDNA repeats, with intermediate-methylation reads excluded from estimates of putatively active rDNA CN (Fig. 6a, right panel). We again detected a strong negative relationship between total rDNA CN, and the proportion of low-methylated rDNA reads (Spearman’s ρ = -0.93; Fig. 6d), indicating that individuals with higher total rDNA CN tend to have a smaller fraction of low-methylated repeats.

We next assessed whether this relationship was sensitive to the thresholds used for methylation calling and read filtering. Base modification calling probabilities showed broadly similar distributions across the 1KGP samples, with two distinct peaks for canonical C and 5mC (Fig. S18a). Restricting calls to P ≥ 0.9 had a larger impact on the detection of unmethylated sites (Fig. S18b). Per-read 5mC distributions generally showed a prominent low-methylation population, frequently concentrated near zero, together with a higher-methylation population of variable prominence across individuals (Fig. S18c). These overall patterns were largely retained after restricting the analysis to high-confidence calls (P ≥ 0.9) (Fig. S18c). The inverse relationship was maintained across multiple filtering strategies (Spearman’s ρ = -0.87 to -0.93) (Figs. S19-20).

These results extend the inverse relationship between total rDNA CN and the proportion of low-methylated repeats to a broader human cohort. Despite substantial inter-individual variation in total rDNA CN and methylation composition, and independently of the thresholds used for site and read methylation detection, individuals with greater numbers of rDNA copies consistently contained a smaller proportion of low-methylated repeats. These findings support the hypothesis that variation in total rDNA CN is partially compensated by differences in the proportion of low-methylated, transcriptionally active/competent repeats, resulting in a more constrained pool of putatively active rDNA copies across individuals.

## Discussion

Interest in rDNA copy number variation (CNV) has grown substantially as altered rDNA dosage has been associated with ageing, developmental disorders, neurodegeneration, and cancer (Rodriguez-Algarra, Whittaker, del Castillo del Rio, & Rakyan, 2025). However, progress in understanding these associations has been hindered by the technical challenge of accurately quantifying rDNA CN and by the difficulty of integrating structural and epigenetic information from the same samples. Here, we described RICO, a computational framework that simultaneously estimates rDNA CN and methylation from Nanopore whole-genome sequencing data, enabling assessment of both total rDNA abundance and the transcriptional competence of individual repeats from a single assay.

By leveraging long reads that span complete rDNA units, RICO overcomes several limitations that have historically complicated rDNA CN estimation in short-read sequencing studies (Hall et al., 2021). Validation across simulated datasets, reference samples and biological systems with established alterations in rDNA regulation demonstrated that RICO yields reproducible CN estimates while simultaneously providing information on rDNA methylation state, confirming the robustness and broad applicability of RICO for studying rDNA CNV.

The main biological insight enabled by our integrated analysis was the consistent inverse relationship between total rDNA CN and the proportion of low-methylated rDNA repeats. This pattern was independently observed across both the GIAB and 1000 Genomes cohorts, suggesting that variation in total rDNA dosage is accompanied by compensatory changes in the fraction of transcriptionally active/competent repeats. As a result, inferred active rDNA CN varied over a substantially narrower range than total CN. While these observations do not demonstrate a defined cellular requirement for a specific number of active rDNA copies, they are consistent with the hypothesis that epigenetic regulation buffers naturally occurring variation in rDNA CN and may help maintain a relatively stable pool of transcriptionally active/competent repeats to ensure sufficient ribosome synthesis and consequently protein translation, while silencing “in excess” rDNA copies to maintain genome stability.

This observation extends and strengthens conclusions from recent long-read studies of human rDNA reporting that individuals with higher rDNA CN tend to harbour a larger fraction of methylated rDNA repeats, resulting in a relatively constrained number of unmethylated copies available for transcription (Hori et al., 2021). Our analyses recapitulate this relationship in two independent cohorts using an orthogonal long-read CN estimation strategy, providing independent support for the idea that epigenetic regulation contributes to the functional buffering of rDNA dosage variation. More broadly, it has been recently emphasized that rDNA CN, chromatin state and transcriptional activity should be considered together rather than as independent properties of rDNA arrays (Hori et al., 2023; Rodriguez-Algarra et al., 2025). Our results support this view by showing that knowledge of total rDNA CN alone provides an incomplete picture of the potentially transcriptionally active/competent rDNA pool.

This model is compatible with emerging views that only a subset of rDNA repeats is required for ongoing transcription, while the remaining repeats may serve structural and regulatory functions (Hori et al., 2023). Having excess rDNA copies may provide a fail-safe architecture that ensures capacity for ribosome biogenesis under cellular demands. Silenced rDNA repeats may also have structural roles in nuclear organization. Heterochromatic rDNA contributes to nucleolar and perinucleolar chromatin organisation and may influence the maintenance of heterochromatin in other genomic regions. Under this model, individuals with greater total rDNA CN could accommodate a larger proportion of inactive repeats while maintaining sufficient transcriptionally active/competent copies for ribosome biogenesis. It is also conceivable that silencing preferentially occurs in damaged or non-functional rDNA repeats to preserve the functional integrity of the rDNA pool. However, whether rDNA methylation is associated with sequence integrity needs to be investigated. Conversely, individuals with fewer copies may require a greater proportion of their rDNA repeats to remain transcriptionally active/competent. Such a mechanism could help explain why substantial variation in total rDNA CN is often tolerated without obvious phenotypic consequences, given that rDNA transcription and ribosome biogenesis are essential in cellular function in all cells. This may also explain why disease associations based on total CN solely have frequently been difficult to reconcile across studies. Whether this relationship observed here reflects active homeostatic regulation or emerges from broader properties of rDNA chromatin organisation remains an important question for future investigation.

Several limitations should be considered. First, rDNA methylation is a proxy for transcriptional competence rather than a direct measure of transcriptional activity. The transcriptional state of intermediately methylated repeats remains particularly uncertain. These repeats may be transcriptionally active/competent or silenced, or potentially represent a dynamic transitional methylation state, and therefore cannot be confidently assigned on the basis of their overall 5mC proportion alone. Moreover, methylation at specific regulatory CpG sites, particularly within the rDNA promoter, may be more directly relevant to transcriptional regulation than the average methylation level across the 45S rDNA locus. For this reason, intermediate-methylated reads were excluded from our estimates of putatively active rDNA CN, and methylation-derived estimates of active CN should be interpreted as relative rather than absolute measures. This issue is particularly evident in the Eμ-*Myc* model, where the relative enrichment of low-methylated repeats in malignant cells was directionally consistent with previous measurements of accessible rDNA chromatin (Diesch et al., 2019), but where estimates of putatively active CN were sensitive to methylation classification thresholds and calling probability filtering.

Another element to consider is that rRNA output is determined not only by the number of transcriptionally active/competent repeats but also by regulating the rate of RNA Polymerase I activity, a major determinant of ribosome biogenesis that is frequently altered in cancer and other growth-associated states (Hein, Hannan, George, Sanij, & Hannan, 2013). Thus, the number of low-methylation repeats should be viewed as one component of rDNA regulation rather than a direct measure of functional ribosome production. Future analyses resolving methylation at specific regulatory CpG sites, particularly within the rDNA promoter, will be important for determining whether particular methylation patterns provide more informative predictors of transcriptional competence than per-read 5mC proportions. Integrating such site-specific methylation profiles with direct measurements of nascent rRNA synthesis, RNA Polymerase I occupancy and chromatin accessibility would further establish how rDNA methylation states relate to actively transcribed rDNA.

Our findings underscore the challenges of obtaining comparable rDNA CN estimates across studies and sequencing platforms. rDNA CN measurements are sensitive to both experimental and analytical factors, including library preparation, sequencing batch effects, reference selection and the strategy used for coverage normalisation (Hall et al., 2021). Importantly, RICO produced highly consistent rDNA CN estimates across independent sequencing runs of the same samples, indicating that RICO is robust to the run-to-run variation. This represents a key methodological advance, particulary given that sequencing batch effects have been identified as a major source of variability in rDNA CN estimation (Hall et al., 2021). Such reproducibility provides an important foundation for reliable comparisons of rDNA CN across experiments and biological conditions.

The differences observed between long-read and short-read estimates likely reflect these factors, as well as fundamental differences in how repetitive rDNA sequences are represented and quantified. Although our results suggest that long-read sequencing may provide a more robust analytical approach for rDNA CN determination, further benchmarking against independent orthogonal measurements will be required to establish the absolute accuracy in rDNA CN estimates. An additional source of variability in rDNA CN estimation is the reference sequence itself. Differences in rDNA reference assembly can influence read mapping and coverage-based estimates. For example, RICO uses a recent long-read-derived human rDNA reference KY962518.1 (Fan et al., 2022), whereas earlier studies relied on older references such as U13369.1 Hall et al. (2021). Although recent telomere-to-telomere assemblies have substantially improved representation of repetitive genomic regions (Nurk et al., 2022), complete reconstruction of chromosome-specific rDNA arrays remains challenging. RICO currently relies on a single representative of the rDNA repeat rather than chromosome-specific arrays and therefore does not capture potential array-level sequence variation. Future improvements in long-read sequencing and assembly technologies should enable the integration of CN, sequence variation, chromatin state and transcriptional output within rDNA arrays, providing a more comprehensive understanding of rDNA regulation.

In conclusion, RICO enables integrated analysis of rDNA CN and methylation from Nanopore long-read sequencing data, providing a scalable approach for studying rDNA variation across populations, developmental states and disease contexts. The high reproducibility of rDNA CN estimates across independent sequencing runs establishes an important foundation for reliable comparisons of rDNA CN across experiments and biological conditions. Beyond this our findings suggest that epigenetic regulation may partially balance extensive variation in total rDNA dosage, highlighting the importance of considering both CN and chromatin state when evaluating the biological consequences of rDNA variation. RICO provides a scalable framework for investigating how genetic and epigenetic variation within rDNA arrays contributes to genome regulation and disease processes across diverse biological contexts.

## Methods

### Nanopore data processing and basecalling

Raw Nanopore signal files (POD5/FAST5) were basecalled using Dorado (v1.2) with methylation calling enabled. Modified BAM (modBAM) files were generated with base-modification tags (MM/ML) retained to allow downstream extraction of CpG methylation information. Basecalling was performed on a high-performance computing (HPC) cluster using GPU nodes and Dorado’s default model for WGS.

### Construction of rDNA-augmented reference genomes

For human samples, the GRCh38 reference genome and the human rDNA reference sequence KY962518.1 were used. A modified GRCh38 reference was generated by appending the complete rDNA repeat sequence as an additional chromosome. To minimise misalignment to partial rDNA-like regions elsewhere in the genome, four highly homologous loci on chromosome 21 (>99% identity to the rDNA unit) were hard-masked, following Fan et al. (2022). To evaluate the effect of tandem repeat context on read alignment and rDNA CN estimation, additional reference genomes containing between one and ten concatenated rDNA repeat units (rDNA×N) as an additional chromosome were constructed. Since Nanopore reads frequently span complete rDNA repeat units, and even multiple consecutive repeats, these augmented references provide sufficient mapping space for ultra-long reads while enabling systematic evaluation of the influence of reference repeat number on alignment behaviour and CN estimation.

For mouse samples, the GRCm39 reference genome and the mouse rDNA reference BK000964.3 were combined as for the human reference. All reference genomes were indexed using minimap2 with the *-d* option. Coordinates for rDNA subregions (5ʹ/3ʹ ETS, 18S, 5.8S, 28S rRNA, ITS1/2, and IGS) for both human and mouse references are provided in Supplementary Table S2 and S3, respectively. Because the mouse rDNA reference is annotated from the 3ʹ end of the repeat unit, the mouse sequence was inverted to position the promoter region at the start of the reference for consistency with the human rDNA reference organisation.

### Read alignment and filtering

Basecalled reads in unaligned modBAM format were converted to FASTQ using samtools fastq, retaining modification tags with parameters: -T “MM,ML”, and aligned with minimap2 (v2.28) to the appropriate rDNA-augmented reference genome withthe parameters: ax map-ont -- secondary=no -Y -y. Aligned reads were sorted and indexed using samtools (v1.22). Reads mapping exclusively to the rDNA chromosome were isolated for the downstream coverage quantification using samtools view.

For CN estimation, coverage across each individual rDNA unit within the augmented reference was quantified independently and subsequently summed to represent total rDNA coverage. This approach prevents artificial dilution of coverage estimates when additional rDNA repeat units are included in the reference genome and ensures that CN estimates are independent of the selected reference configuration. Following alignment, only primary alignments were retained to minimise contributions from residual rDNA-like regions elsewhere in the genome. The resulting aligned modBAM file formed the input for both CN estimation and methylation analysis.

### Mean-based CN estimation

RICO estimates rDNA CN using a mean-coverage approach in which sequencing coverage across the complete 45S rDNA transcription unit is normalised against the coverage of curated SCG reference panels. For benchmarking purposes, a previously described mode-based strategy (Sharma et al., 2022) was additionally implemented for comparison.

Coverage across the complete 45S rDNA transcription unit (5ʹETS-28S) was computed using bedtools coverage, with genomic intervals defined by an rDNA BED annotation corresponding to the selected rDNA×N reference. The quantified region comprised the 5ʹ external transcribed spacer (5ʹETS), 18S, internal transcribed spacer 1 (ITS1), 5.8S, internal transcribed spacer 2 (ITS2), 28S, and 3ʹ external transcribed spacer (3ʹETS).

For augmented references containing multiple concatenated rDNA repeat units (rDNA×N), coverage was calculated independently for the 45S region of each repeat and subsequently summed to obtain the total rDNA coverage. This prevents artificial dilution of coverage estimates resulting from increasing the number of rDNA repeat units in the reference genome and ensures that rDNA CN estimates are independent of the selected reference configuration.

For background normalisation, sequencing coverage across the SCG panels (SCG-1, SCG-2 and SCG-3 for human samples; mSCG for mouse samples) was calculated using samtools depth. Mean SCG coverage was calculated as the total sequencing depth divided by the number of genes within the corresponding SCG panel. rDNA CN was then calculated as:

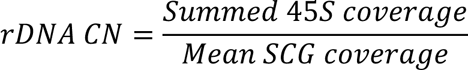

### Mode-based CN estimation

The alternative mode-based strategy was implemented according to Sharma et al. (2022). Briefly, sequencing coverage was calculated using 1 kb sliding windows across the rDNA locus, and rDNA CN was estimated from the modal coverage value after normalisation to the corresponding SCG panel. This implementation was used exclusively for benchmarking against the mean-coverage approach.

In this approach, per-based read depth across both single-copy genes and rDNA is first extracted using *Samtools*. Using a 1000 bp sliding window, the mean coverage is then computed, and these windowed coverage values are binned. After that, the top three most frequent coverage bins, i.e. the top three modes for both the single-copy gene panel and rDNA region are identified. These six values are then used to generate a matrix table. The resulting 36 CN values are calculated by dividing rDNA coverage by the background coverage in each pairwise combination from the matrix. The final CN estimate is the average of these 36 values in the matrix.

### rDNA methylation profiling and classification

RICO performs DNA methylation quantification from aligned modBAM using *Modkit* and generates two complementary methylation outputs: per-site CpG methylation profiles and per-read CpG methylation profiles. Per-site profiles describe methylation at individual CpG positions across the rDNA repeat, whereas per-read profiles retain CpG calls associated with individual rDNA reads, enabling methylation states to be examined at the level of individual rDNA repeat units. The analyses presented in this study primarily used the per-read methylation output to characterise heterogeneity in methylation among individual rDNA repeats.

Because Nanopore reads can span multiple rDNA repeat units, reads were divided into individual read/repeat units according to their mapping coordinates (Supplementary Table S2 and S3). CpG calls assigned to any of the repeat units in the augmented mapping reference were then projected onto a common single-repeat coordinate. Thus, each read/repeat unit represents the portion of a sequencing read corresponding to one 45S rDNA repeat unit rather than necessarily one independent sequencing read.

For each rDNA read/repeat unit, canonical cytosine (C) and 5-methylcytosine (5mC) calls at CpG sites were extracted together with their corresponding call probabilities. Analyses were performed either using all available CpG calls (i.e. unfiltered) or retaining only high-confidence calls with a calling probability of *P* ≥ 0.9. The methylation level of each retained read/repeat unit was calculated as the proportion of 5mC calls:

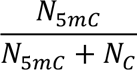

where N_5mC_ and N_C_ represent the numbers of 5mC and canonical C calls, respectively. Read/repeat units were classified as low-methylated (<0.3), intermediate-methylated (0.3-0.7), or high-methylated (>0.7). Alternative thresholds of 0.1 and 0.9 were additionally evaluated to assess the sensitivity of methylation classification to threshold selection.

The robustness of read-level methylation classification was further assessed by varying the minimum number of CpG calls required per read/repeat unit. Analyses were performed with no minimum requirement or with minimum thresholds of ≥10, ≥50, or ≥100 CpG calls per read/repeat unit, in combination with either unfiltered or high-confidence (*P* ≥ 0.9) CpG calls.

To estimate the number of putatively transcriptionally active/competent rDNA copies, the fraction of low-methylated read/repeat units was used as a proxy for the fraction of transcriptionally active/competent rDNA repeats. Putatively active rDNA CN was calculated as total rDNA CN × the fraction of low-methylated read/repeat units. Intermediate-methylated units were excluded from this estimate because their transcriptional state could not be confidently assigned from repeat-wide methylation alone.

For spatial methylation analysis, reads spanning multiple tandem rDNA repeat units were divided into individual read/repeat units based on their mapping coordinates (as described above). Calls assigned to any of the repeat units in the augmented mapping reference were projected onto a common single-repeat coordinate, allowing the spatial distribution of canonical C and 5mC calls to be visualised across individual 45S rDNA repeat units. To minimise the inclusion of partially covered repeat units, i.e. short rDNA fragments and improve visualisation of methylation patterns across the 45S rDNA region, only read/repeat units with a CpG-call span of at least 10 kb were retained. CpG-call span was defined as the distance between the first and last retained CpG calls within each read/repeat unit.

### Construction of SCG sets

#### Human SCG panels

To generate reference panels of putative SCGs as background coverage controls, the GENCODE primary gene annotation (GRCh38.p14) and the Catalogue of Somatic Mutations In Cancer (COSMIC) cancer gene set were obtained. Genes listed in COSMIC were removed to avoid regions known to undergo recurrent somatic alterations, as were genes shorter than 300 bp to minimise coverage instability. Genes located on the sex chromosomes were removed to restrict the panels to loci with consistent diploid dosage across individuals regardless of sex.

To reduce ambiguity arising from paralogous or partially duplicated loci, each candidate gene was aligned against the full gene set using BLAST. Genes with significant sequence similarity to any other gene (E-value < 1×10⁻⁶) were removed. We further refined the panel by re-running BLAST with increasing word-size parameters (100, 500, 1000, 1500, and 2000 bp). Larger word sizes require longer exact-match seeds and therefore preferentially eliminate genes with weaker or fragmented homology, allowing us to progressively filter out loci that might otherwise introduce ambiguous mapping. These filtering steps then produced three preliminary SCG panels corresponding to word-size thresholds (100, 1500, and 2000 bp). Genes located on the sex chromosomes were excluded to restrict the panels to loci with consistent diploid dosage across individuals regardless of sex.

To further improve coverage stability, we applied an additional round of filtering to remove genes likely to be affected by mapping artefacts or repetitive content. First, intronless genes, i.e. single-exon genes were excluded, as exploratory analyses revealed that these loci frequently exhibited highly variable coverage across samples. Second, we removed genes overlapping annotated repetitive elements using RepeatMasker annotations, except for simple repeats and low-complexity repeats, which are less likely to influence coverage estimates.

These filtering steps yielded three progressively refined SCG panels that differed in the number of retained genes: 174 genes in SCG-1, 373 genes in SCG-2, and 514 genes in SCG-3 (corresponding to the 100, 1500, and 2000 bp word-size thresholds, respectively). Despite reductions in panel size, the mean and median coverage across genes remained highly consistent between versions, while the distribution of gene-level coverage became markedly less variable following removal of intronless and repeat-associated loci (add results to Suppl?). These three progressively filtered SCG panels were subsequently benchmarked to evaluate the influence of reference gene selection on rDNA CN estimation.

#### Mouse SCG panels

Mouse SCG panels were constructed using an analogous workflow. Human COSMIC genes were first mapped to mouse orthologues to remove COSMIC-associated loci from the mouse gene set. Following BLAST filtering using word sizes of 1000, 1500, and 2000 bp, intronless genes and repeat-associated loci were removed. This resulted in 44 genes (SCG-1), 141 genes (SCG-2), and 196 genes (SCG-3). Given the limited size of SCG-1 and the similar performance of SCG-2 and SCG-3, a single mouse SCG panel (mSCG, 141 genes) was retained for downstream analyses.

#### Simulated reads

Nanopore reads were simulated using Badread (Wick, 2019), which generates synthetic reads by fragmenting reference DNA, ligating adapters, and introducing substitutions, insertions, deletions, and low-complexity artifacts according to their empirically derived error models. Simulation parameters were selected to reproduce the bimodal read-length distribution observed in two independent PromethION sequencing runs of HG002.

The simulation reference genome was derived from the human GRCh38 assembly and included autosomal chromosomes 1-22 only, as all SCGs are located on autosomes. As in the alignment step for real sequencing data, genomic regions with high sequence similarity to the rDNA reference were masked to minimise ambiguous read alignment (Fan et al., 2022).

To model different rDNA CN scenarios, the complete human reference rDNA sequence (KY962518.1, ∼44 kb) was appended to the reference genome as 100 or 300 concatenated rDNA units, as a single, separate chromosome.

Nanopore reads were simulated from each reference genome using the “pretty good” Badread preset, with a target mean genomic coverage of 40x, calculated based on the total size of the reference genome including the rDNA insert. Default Badread parameters were used unless otherwise specified.

For each rDNA CN scenario (100 and 300 copies), two independent simulations were performed to capture the bimodal read-length characteristics observed in empirical HG002 sample Nanopore data. For the short-read length distribution, a mean fragment length of 900 bp and a standard deviation of 100 bp were used. For the long-read length distribution, a mean fragment length of 20 kb and a standard deviation of 10 kb were set. Reads from these two simulations for each scenario were then combined to form the final benchmarking datasets.

Alignments of the simulated reads were performed directly from FASTQ files rather than modBAM files as these reads do not contain base-modification information. All downstream processing steps, including alignment to rDNA-augmented references, filtering of primary alignments, and rDNA CN estimation using both mean- and mode-based coverage approaches, were identical to those applied used for the real Nanopore sequencing data.

#### Genome in a Bottle (GIAB) trio samples

Publicly available Oxford Nanopore whole-genome sequencing datasets were obtained for six Genome in a Bottle (GIAB) samples derived from lymphoblastoid cell lines, comprising the Ashkenazi trio (HG002, HG003 and HG004) and Han Chinese trio (HG005, HG006 and HG007). For each individual, data from two independent PromethION R10.4.1 flow cells (FLO-PRO114M), generated using the Ligation Sequencing Kit V14 chemistry (SQK-LSK114), were analysed as technical replicates to assess reproducibility of rDNA CN estimation and methylation profiling. Dataset descriptions, sample metadata, and sequencing summaries are provided within the corresponding ONT GIAB S3 resources.

In addition, we generated an independent in-house WGS dataset for HG002 using the same PromethION FLO-PRO114M flow-cell chemistry and SQK-LSK114 library preparation protocol. This dataset was processed alongside the publicly available GIAB datasets as an additional assessment of reproducibility.

#### Mouse ESC samples

WT and *Atrx*-KO mouse ESCs have been previously described, including their derivation and culture conditions, by Udugama et al. (2018). HMW genomic DNA was extracted from WT and Atrx KO ESCs using the Monarch® HMW DNA Extraction Kit (New England Biolabs), following the manufacturer’s instructions with conditions optimised to preserve long DNA fragments. Cells were harvested at similar densities to minimise confounding effects of cell state on DNA quality, and DNA preparations were assessed for concentration, purity, and fragment size before library preparation.

DNA was then sheared using a Megaruptor system, followed by barcoded Nanopore library preparation using the Oxford Nanopore Native Barcoding Kit (SQK-NBD114-24). WT and KO samples were multiplexed and sequenced together on a single Oxford Nanopore PromethION R10.4.1 flow cell (FLO-PRO114M). Flow-cell washes and reloads were performed where possible during sequencing to maximise data yield.

#### Eμ*-Myc* mouse lymphoma samples

All animal experiments were conducted in compliance with the ethics protocol approved by the Australian National University’s Animal Experimentation Ethics Committee project A2018/46. A heterozygous Eμ-*Myc* colony was maintained by mating transgenic positive males of C57BL/6J background with wild-type C57BL/6J females. Wild-type mice aged 4-10 weeks and transgenic positive mice aged 10-25 weeks exhibiting clear signs of lymphoma, such as enlarged lymph nodes, rapid breathing, hunching, slow movement and ruffled fur, were sacrificed for sample collection. Lymphomas arising in this model can originate from either IgM-negative (IgM-) or IgM-positive (IgM+) clone. To reduce potential confounding variability, we used the more prevalent pre-B cell population (B220+/IgM-) in this study.

Pre-B cells were isolated from bone marrow and sorted via Fluorescence-Activated Cell Sorting (FACS). Genomic DNA was extracted from 10^6^ sorted pre-B cells per samples using the Monarch HMW DNA Extraction Kit for Cells & Blood (NEB Cat. No. #T3050) according to the manufacturer’s instructions.

HMW genomic DNA libraries were prepared using the Oxford Nanopore ultra-long DNA sequencing kit (SQK-ULK114) and sequenced on Oxford Nanopore PromethION R10.4.1 flow cells (FLO-PRO114M). The WT sample was sequenced on a single PromethION flow cell. The MAL sample was initially sequenced on one flow cell, however, reduced pore performance and library overloading were observed during sequencing. Therefore, the same MAL library was reloaded onto a second flow cell at approximately half of the original loading concentration. Sequencing data from both MAL runs were subsequently combined for downstream analyses.

#### Human lymphoblastoid cell lines

Both human lymphoblastoid cell lines, HG00326 and HG03069 were purchased form Coriell Institute for Medical Research. Both cell lines were maintained in RPMI-1640 supplemented with 15% Fetal Bovine Serum (FBS) and GlutaMax (Thermo). Prior to DNA extraction cells were seeded at 0.6 x10^6^ cells per ml in 3 ml complete media in a 6 well plate and incubate overnight. Genomic DNA was extracted from 2 x10^6^cells per samples using the Monarch HMW DNA Extraction Kit for Cells & Blood (NEB Cat. No. #T3050) according to the manufacturer’s instructions for long read sequencing or the Macherey-Nagel NucleoSpin Tissue DNA, RNA and protein isolation kit (MN cat. No. 740952.25) for qPCR analysis. DNA concentrations for qPCR analysis were measured using the Nanodrop.

For Nanopore WGS, barcoded libraries were prepared using the Oxford Nanopore Native Barcoding Kit (SQK-NBD114-24). HG00326 and HG03069 were multiplexed and sequenced together on a single PromethION R10.4.1 flow cell (FLO-PRO114M). Flow-cell washes and library reloads were performed where possible during sequencing to maximise data yield.

#### Copy number qPCR

QPCR analysis was performed using 20ng of gDNA per reaction on the StepOne Plus Real-Time PCR System (Applied Biosystems) using a SYBRGreen Fast Master Mix (Applied Biosystems) with the following thermocycling conditions: denaturation for 10 s at 95 °C and amplification for 30 s at 61 °C for a total of 40 cycles. Primers pairs (Merck) specific for the 28S, 18S, IGS rDNA region and two genes expected to be present at a stable single CN in these lymphoblastoid cell lines and are independent of the rDNA locus, *TP53* and *RPPH1* (Table 1) were used. For each replicate relative 28S, 18S and IGS Ct values were normalised to single copy gene *TP53* or *RPPH1* Ct values (delta CT) and fold change over HG00326 was calculated via the delta delta Ct method. Dissociation curves for each sample were analysed to control for unspecific amplification.

**Table 1.**
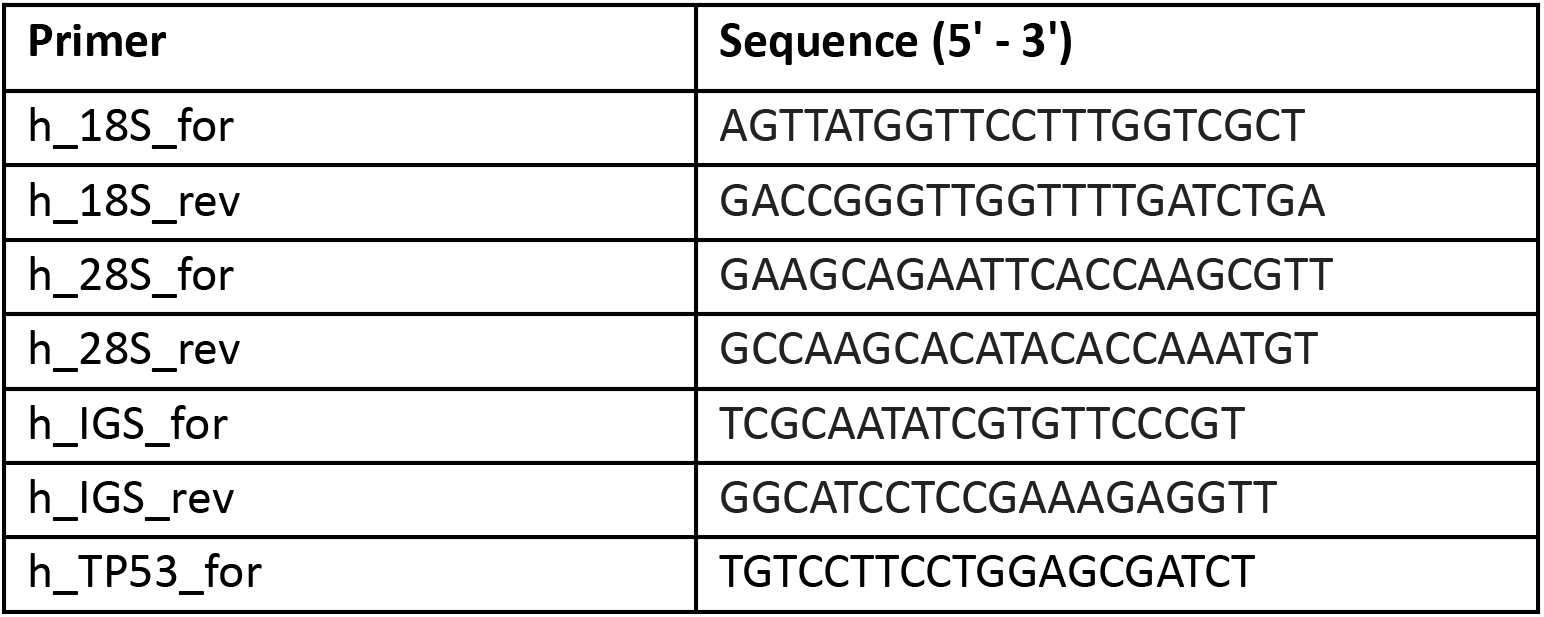
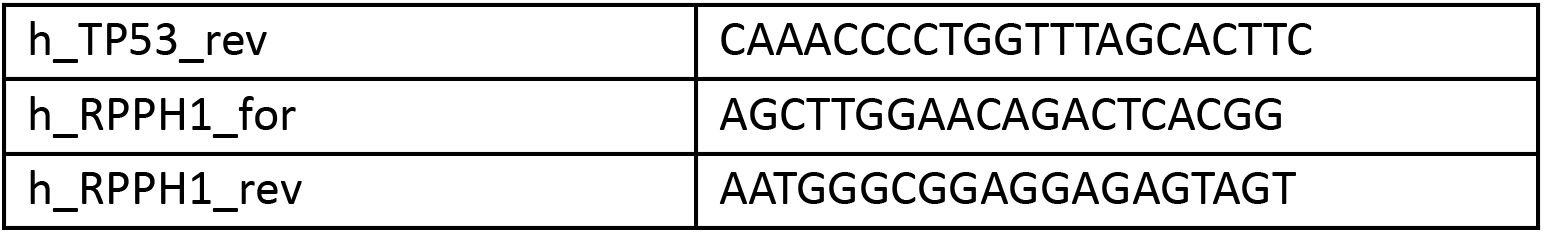
Primer sequences used in qPCR.

#### 1000 Genomes Project (1KGP) samples

Previously published short-read rDNA CN estimates for 1KGP samples were obtained from (Hall et al., 2021). Briefly, Hall et al. aligned short-read WGS data using BWA-MEM and estimated CN from average sequencing depth across defined regions of the 18S and 28S rRNA genes, normalised to average chromosome 1 sequencing depth. The analysed regions corresponded to positions 3841-3985 (18S) and 8049-8198 (28S) of the human 45S rDNA reference sequence U13369.1.

The 1KGP-ONT samples analysed in this study were prepared using ONT sequencing kit (SQK-LSK114) and sequenced on Oxford Nanopore PromethION R10.4.1 flow cells (FLO-PRO114M).

## Data Availability

Oxford Nanopore WGS data for the GIAB Ashkenazi trio (HG002, HG003, HG004) and Han Chinese trio (HG005, HG006, HG007) were obtained from publicly available ONT reference datasets (GIAB 2023.05 and 2025.01 releases). Dataset descriptions are available at: https://epi2me.nanoporetech.com/giab-2023.05/ and https://epi2me.nanoporetech.com/giab-2025.01/. Sequencing data are hosted in the AWS S3 repositories: s3://ont-open-data/giab_2023.05/ and s3://ont-open-data/giab_2025.01/.

Oxford Nanopore WGS datasets for 1KGP samples used in this study were generated by the 1KGP Long-Read Sequencing Consortium, using high-coverage PromethION sequencing of high-molecular-weight DNA derived from Coriell 1KGP cell lines. Details regarding the consortium and sequencing effort are available through the consortium resource (https://millerlaboratory.com/1KGP-LRS.html) and associated publication by Gustafson et al. (2024). The datasets analysed in this study were obtained through the ONT-1KGP resource hosted at the Australian National Computational Infrastructure (NCI), Australia, under NCI project de95 (https://geonetwork.nci.org.au/geonetwork/srv/eng/catalog.search#/metadata/f6758_587 <u>1_3955_7179</u>). This resource was established through collaboration between Zaka Yuen and Hasindu Gamaarachchi to provide Australian researchers with local HPC-scale access to long-read population sequencing datasets, including raw POD5/BLOW5 signal data and processed Nanopore sequencing outputs.

Oxford Nanopore WGS datasets generated in this work (GIAB and HG002 samples, WT and *Atrx*-KO mESC samples, and Eμ-*Myc* lymphoma samples) are available at the NCBI Sequence Read Archive (SRA) under BioProject accession PRJNA1473546.

RICO source code and documentation are available at: https://github.com/comprna/RICO.

## Author Contributions

EE, NH and ZY conceptualised the study. ZY led the software development, methodology, validation and computational analyses under the supervision of EE and NH. NL contributed to the analysis of the 1KGP datasets. TU generated the *Eμ-Myc* mouse models, AG performed mESC sample preparation, and LW provided mESC biological samples. ZY led data curation and visualisation and wrote the original manuscript, with contributions from EE and NH. AG and RH provided critical feedback and contributed substantially to the manuscript review. All authors reviewed and approved the manuscript.

## Acknowledgements

We thank Hasindu Gamaarachchi (UNSW Sydney) for collaborating in the preparation of the 1KGP samples at the National Computing Infrastructure (NCI).

This research was supported by the Australian Research Council (ARC) Discovery Project grants DP220101352, DP250100070, and DP250103133; by the National Health and Medical Research Council (NHMRC) Ideas Grant 2018833; and by a grant from the New Zealand Marsden Fund (14-MAU-053). This research was also indirectly supported by the Australian Government’s National Collaborative Research Infrastructure Strategy (NCRIS) through access to computational resources provided by the National Computational Infrastructure (NCI) through the National Computational Merit Allocation Scheme (NCMAS) and the ANU Merit Allocation Scheme (ANUMAS). The funding bodies had no role in study design, data collection, or data analysis.

## Supplementary Figures

**Supplementary Figure S1.**
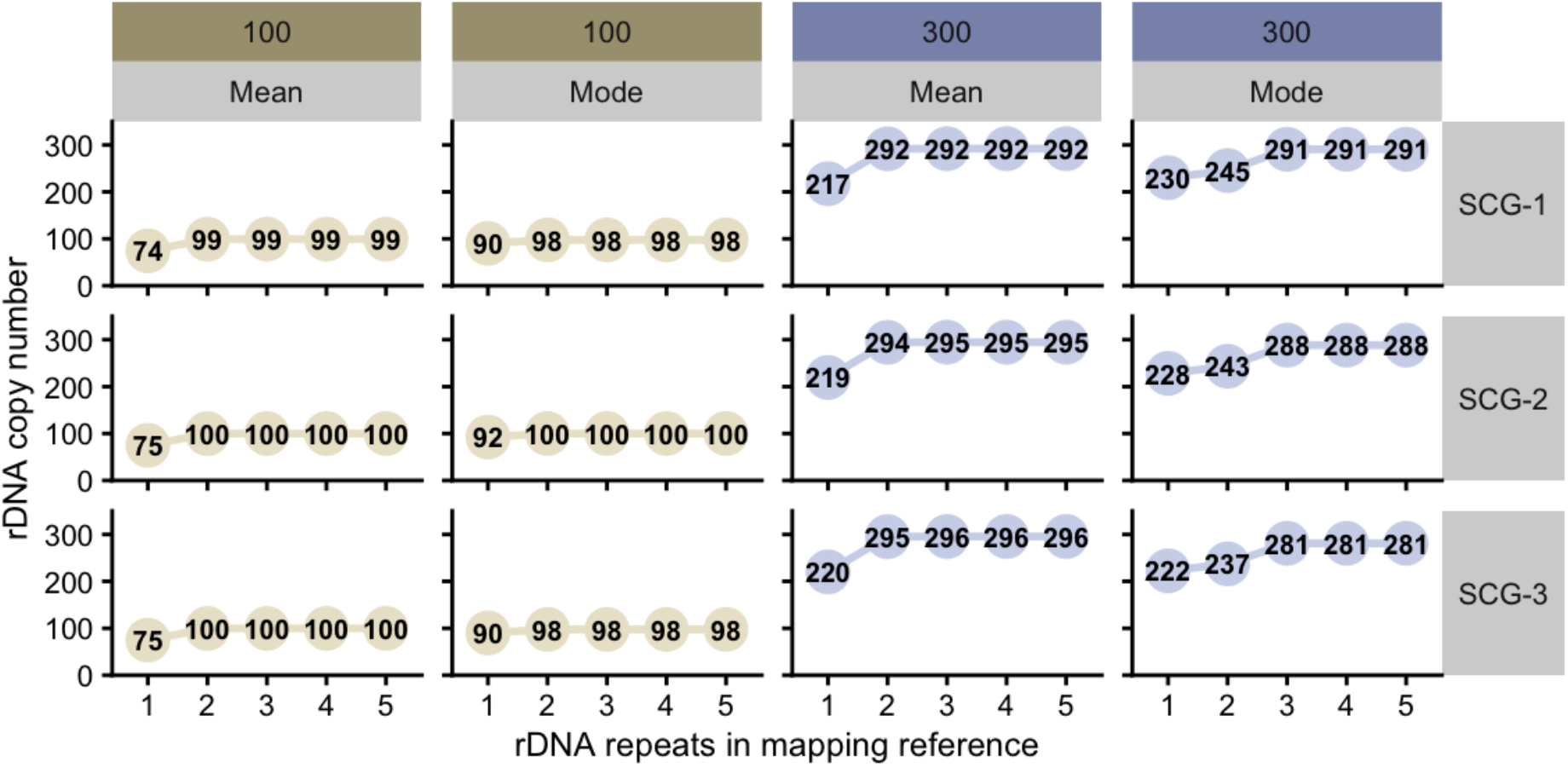
Comparison of mean-coverage and mode-coverage approaches for rDNA copy number (CN) estimation in simulated datasets. Estimated rDNA CN across different numbers of rDNA repeats included in the mapping reference for simulated datasets containing 100 (left) or 300 (right) known rDNA copies. CN estimates were calculated using either the mean-coverage or mode-coverage approach and normalised using three different single-copy gene (SCG) reference panels (SCG-1, SCG-2, SCG-3).

**Supplementary Figure S2.**
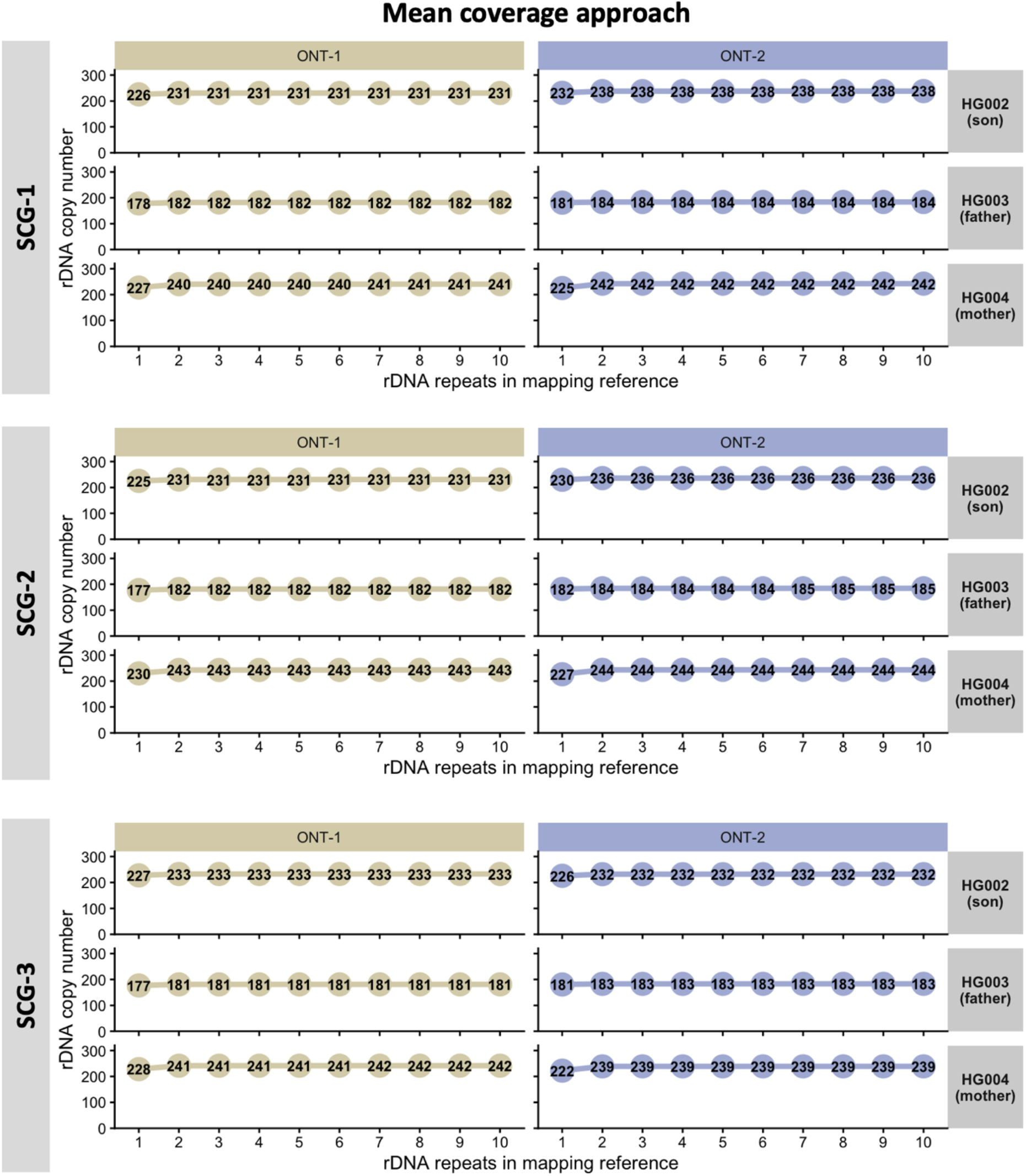
rDNA CN estimation using the mean coverage approach in Ashkenazi Trio samples. Estimated rDNA copy number (CN) (y-axis) as a function of the number of rDNA repeats included in the mapping reference (x-axis; 1–10 repeats) for the Ashkenazi trio samples HG002 (son), HG003 (father), and HG004 (mother). Results are shown for two independent Oxford Nanopore sequencing datasets (ONT-1 - left and ONT-2 - right) and for three single-copy gene reference (SCG) panels (SCG-1 - top; SCG-2 - middle; SCG-3 - bottom).

**Supplementary Figure S3.**
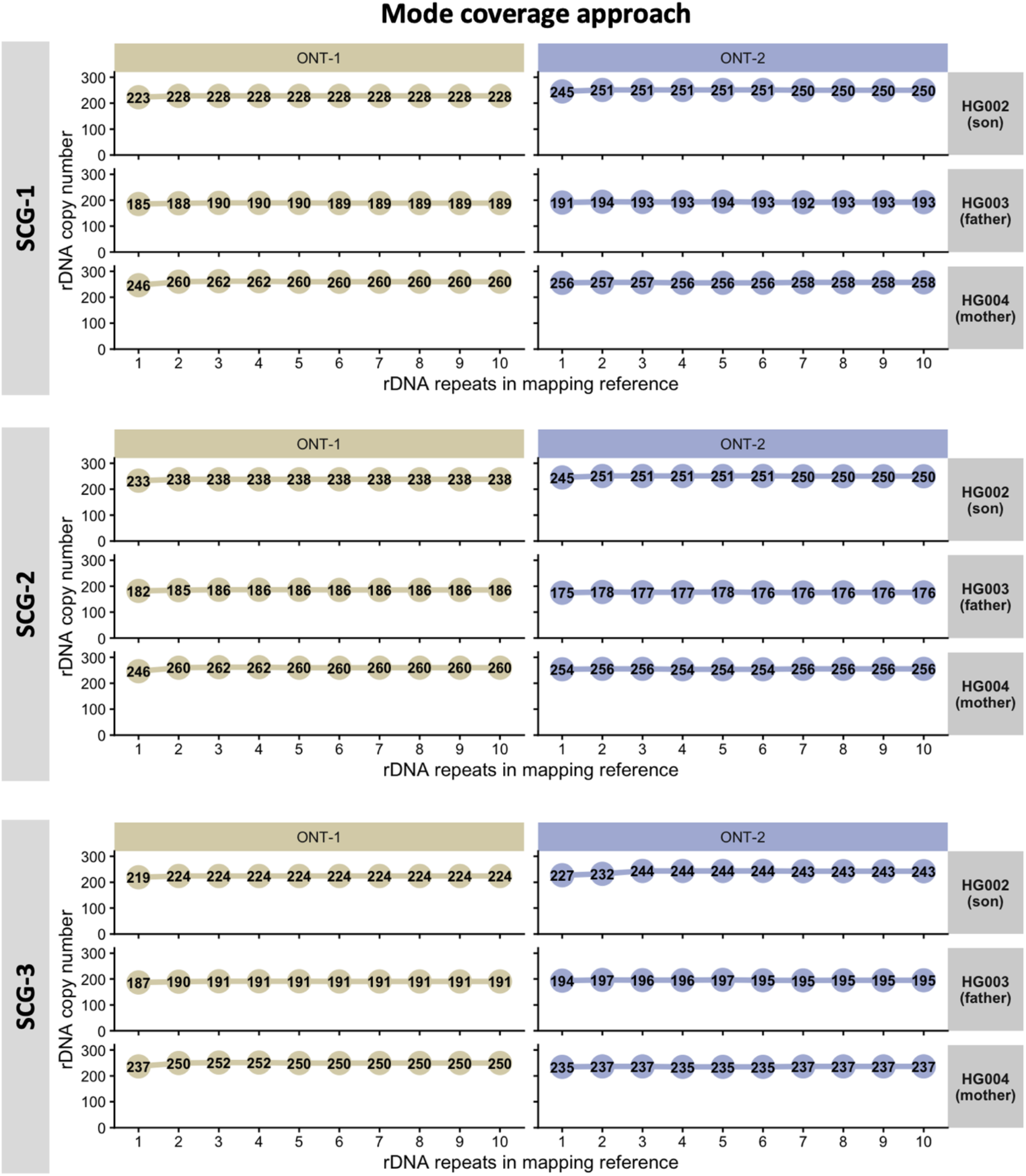
rDNA CN estimation using the mode coverage approach in Ashkenazi Trio samples. Estimated rDNA copy number (CN) (y-axis) as a function of the number of rDNA repeats included in the mapping reference (x-axis; 1–10 repeats) for the Ashkenazi trio samples HG002 (son), HG003 (father), and HG004 (mother). Results are shown for two independent Oxford Nanopore sequencing datasets (ONT-1 - left and ONT-2 - right) and for two single-copy gene (SCG) reference panels (SCG-1 - top; SCG-2 - middle; SCG-3 - bottom).

**Supplementary Figure S4.**
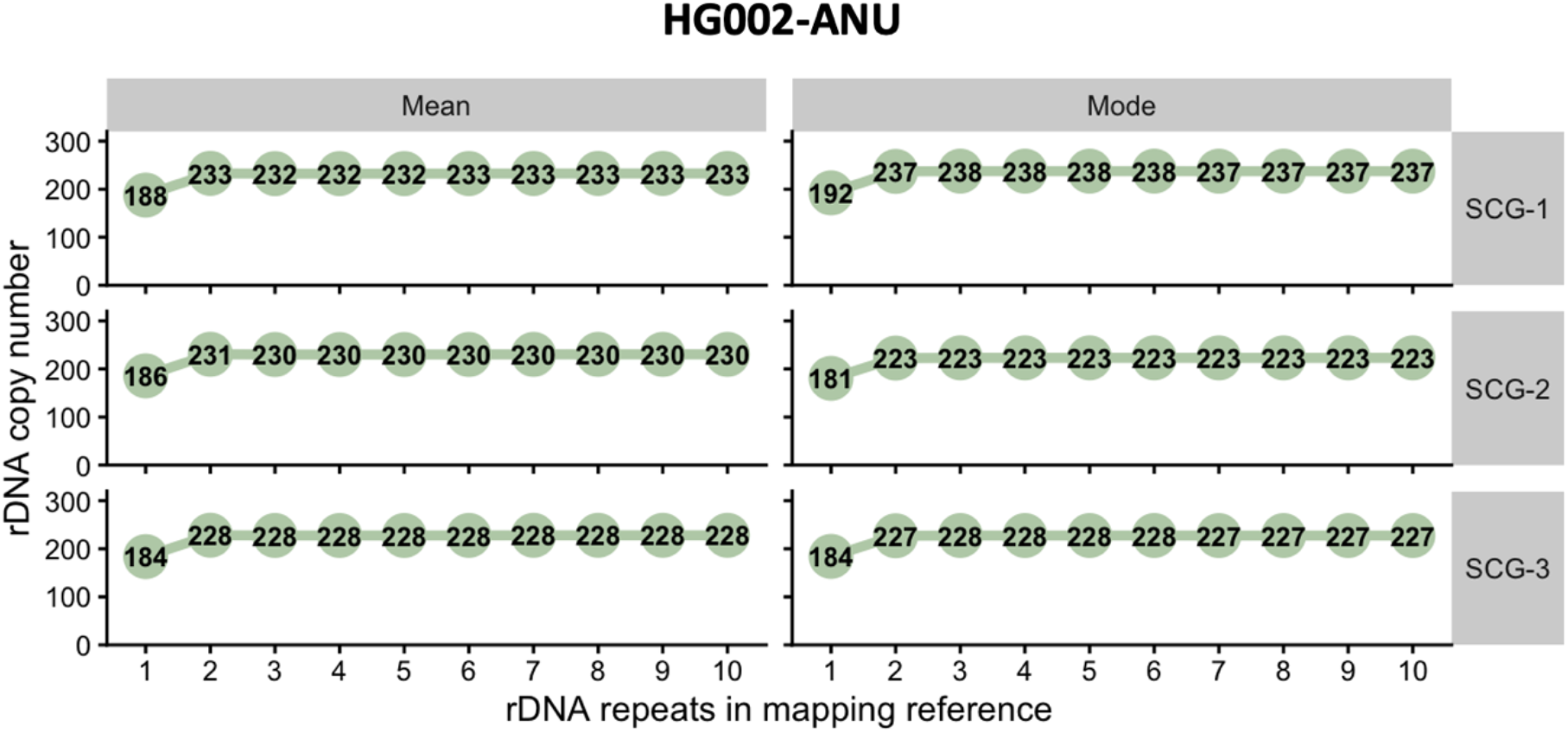
rDNA CN estimation for the in-house HG002 sample (HG002-ANU) using mean- and mode-coverage approaches across three SCG panels. Estimated rDNA copy number (CN) (y-axis) as a function of the number of rDNA repeats included in the mapping reference (x-axis; 1–10 repeats) for HG002-ANU. Results are shown for two coverage-based estimation methods (mean, left; mode, right) and for three single-copy gene (SCG) reference panels (SCG-1 - top; SCG-2 - middle; SCG-3 - bottom).

**Supplementary Figure S5.**
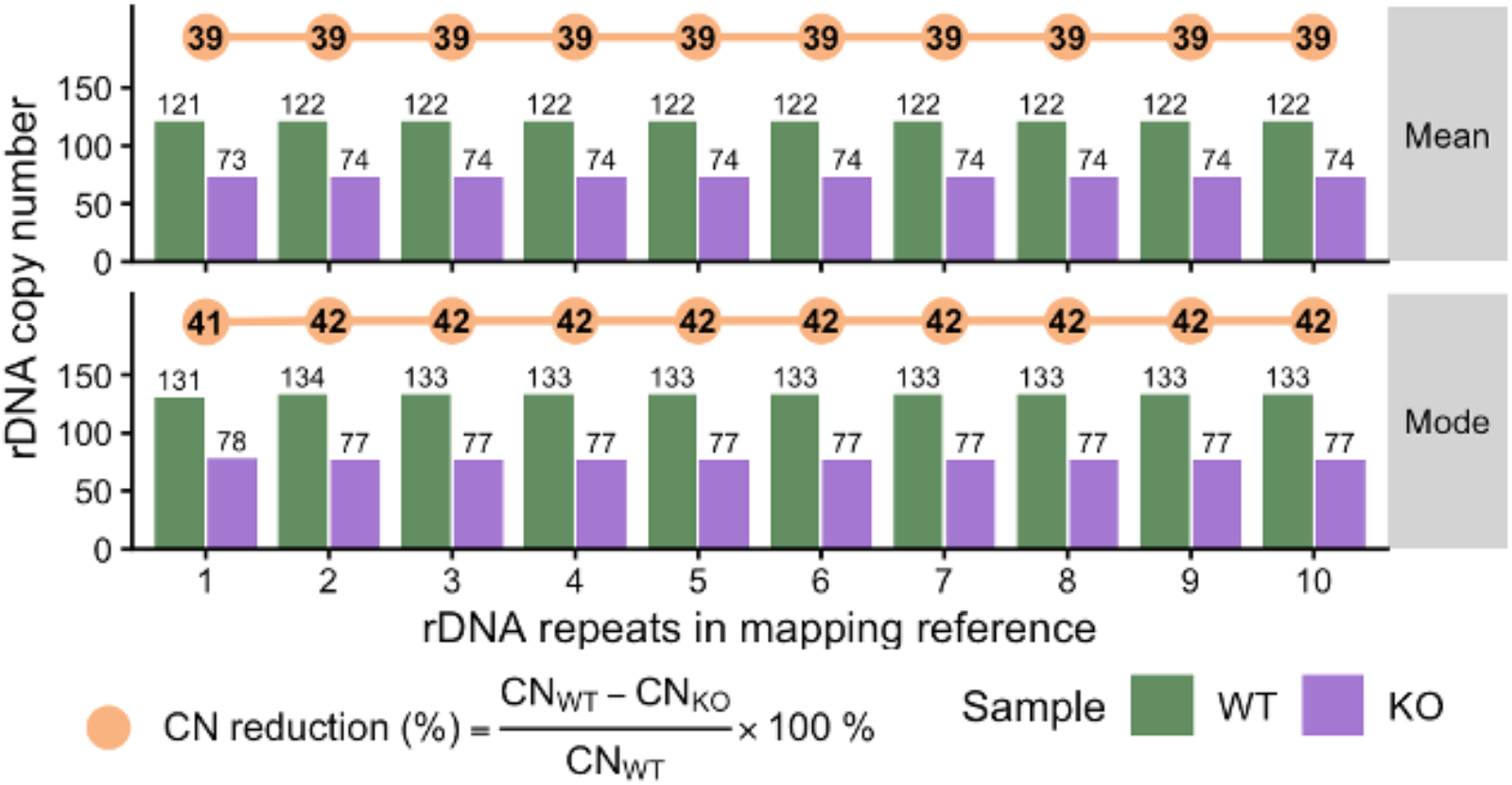
rDNA copy-number (CN) estimates in *Atrx*-KO mouse embryonic stem cells (mESCs) are stable across mapping reference configurations. Estimated rDNA CN for wild-type (WT) and *Atrx*-KO mESCs across augmented mapping references containing 1-10 rDNA repeats. Bars show CN estimates obtained using the mean-coverage (top) and mode-coverage (bottom) approaches. Orange markers indicate the percentage reduction in rDNA CN in *Atrx*-KO relative to WT cells.

**Supplementary Figure S6.**
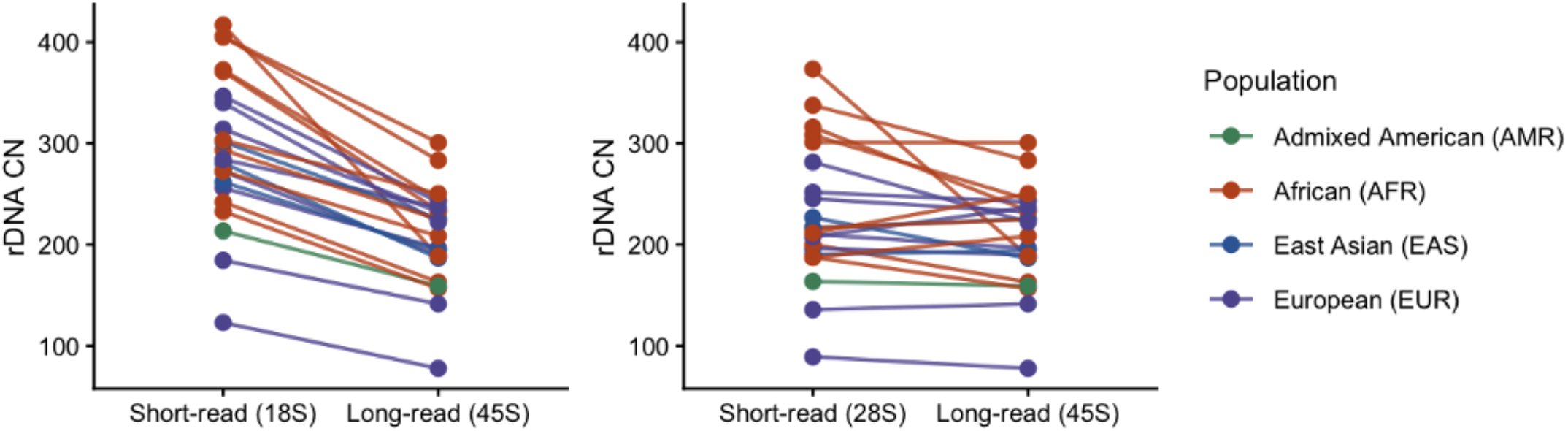
short-read vs long-read sequencing approaches. Comparison of rDNA CN estimates obtained from short-read sequencing based on the 18S rRNA region (left) or the 28S rRNA region (right) and long-read Nanopore sequencing based on the full 45S rDNA transcription unit. Each point represents an individual sample, with paired estimates from short-read and long-read data connected by lines. Samples are coloured by continental ancestry.

**Supplementary Figure S7.**
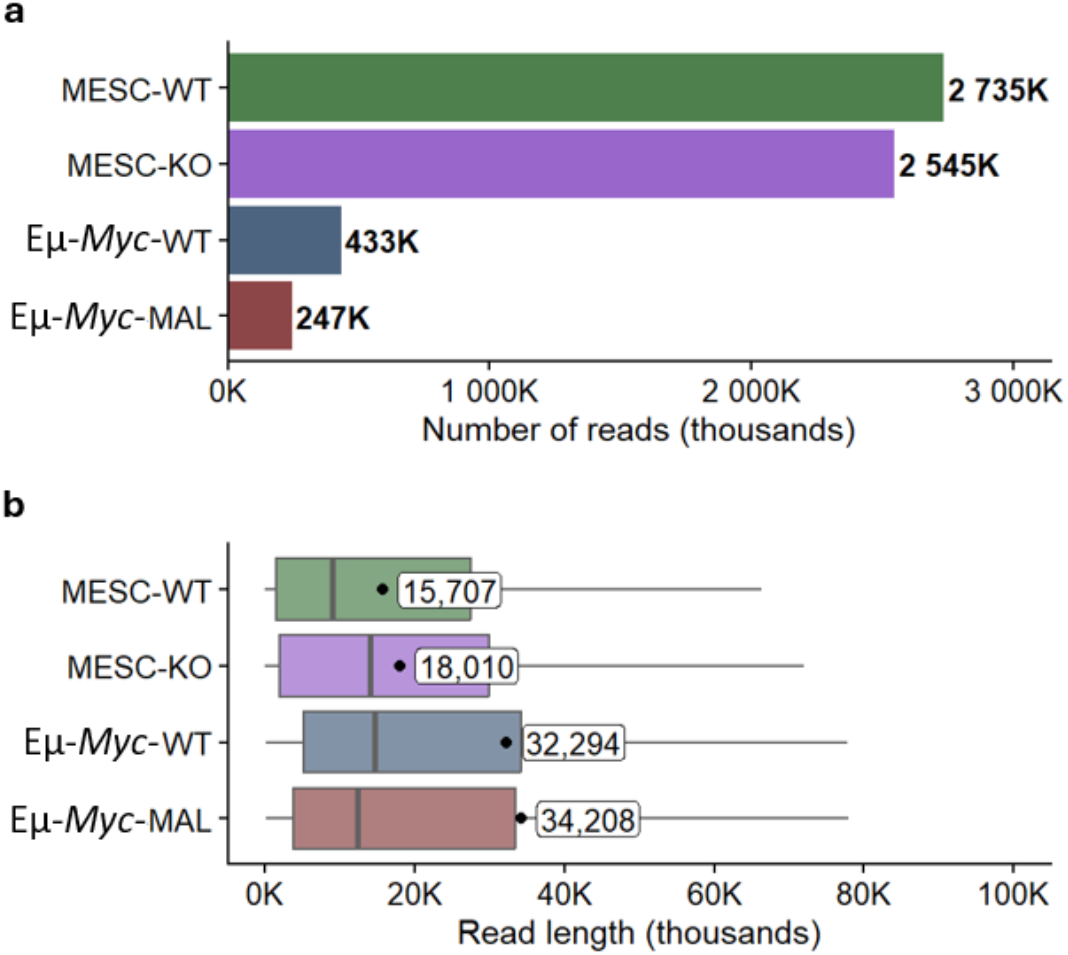
Sequencing statistics for mouse samples. **(a)** Total number of reads generated for each mouse sequencing dataset, obtained from the unaligned modified BAM (modBAM) files. Values shown to the right of each bar indicate the total number of reads (in millions) per run. **(b)** Read-length distributions for each mouse sequencing dataset, derived from unaligned modBAM files. Boxplots summarise the read length distribution, with black dots and accompanying labels indicate the mean read length for each run.

**Supplementary Figure S8.**
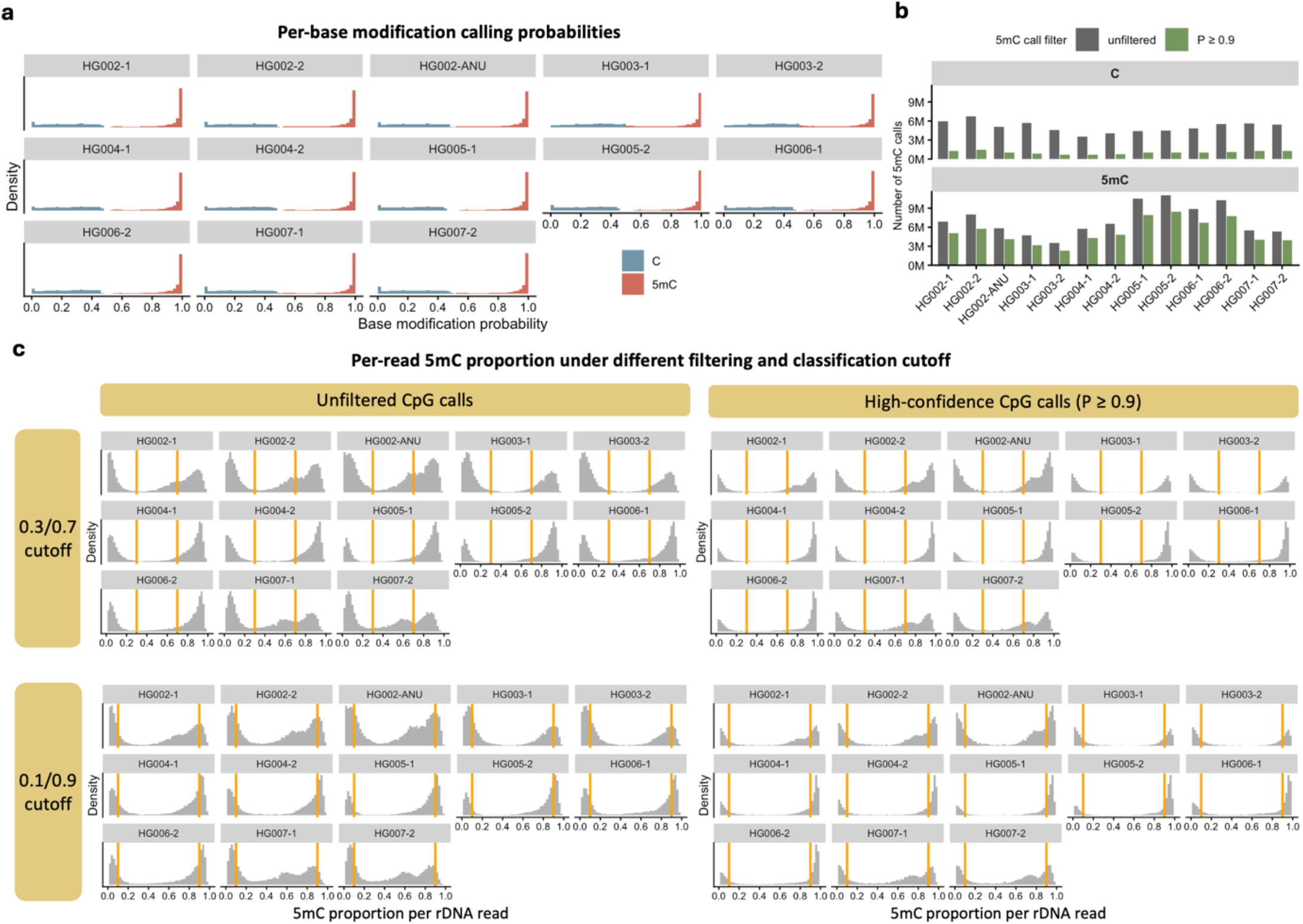
Evaluation of 5mC calling probability filtering and read-level methylation classification thresholds. **(a)** Distribution of per-base modification calling probabilities (P) for canonical C and 5mC calls across rDNA reads from the trio samples (HG002-HG007). For visualization on a common modification-probability scale, canonical C calls are plotted as 1 − P, whereas 5mC calls are plotted as P. **(b)** Number of canonical C and 5mC calls before filtering (unfiltered) and after retaining high-confidence calls with P ≥ 0.9. Application of the probability threshold preferentially reduces canonical C calls, reflecting differences in the calling-probability distributions of canonical and modified CpGs. **(c)** Distribution of the per-read 5mC proportion calculated using either unfiltered CpG calls (left) or high-confidence CpG calls with P ≥ 0.9 (right). Alternative read-level methylation classification thresholds of 0.3/0.7 (top) and 0.1/0.9 (bottom) are indicated by orange vertical lines.

**Supplementary Figure S9.**
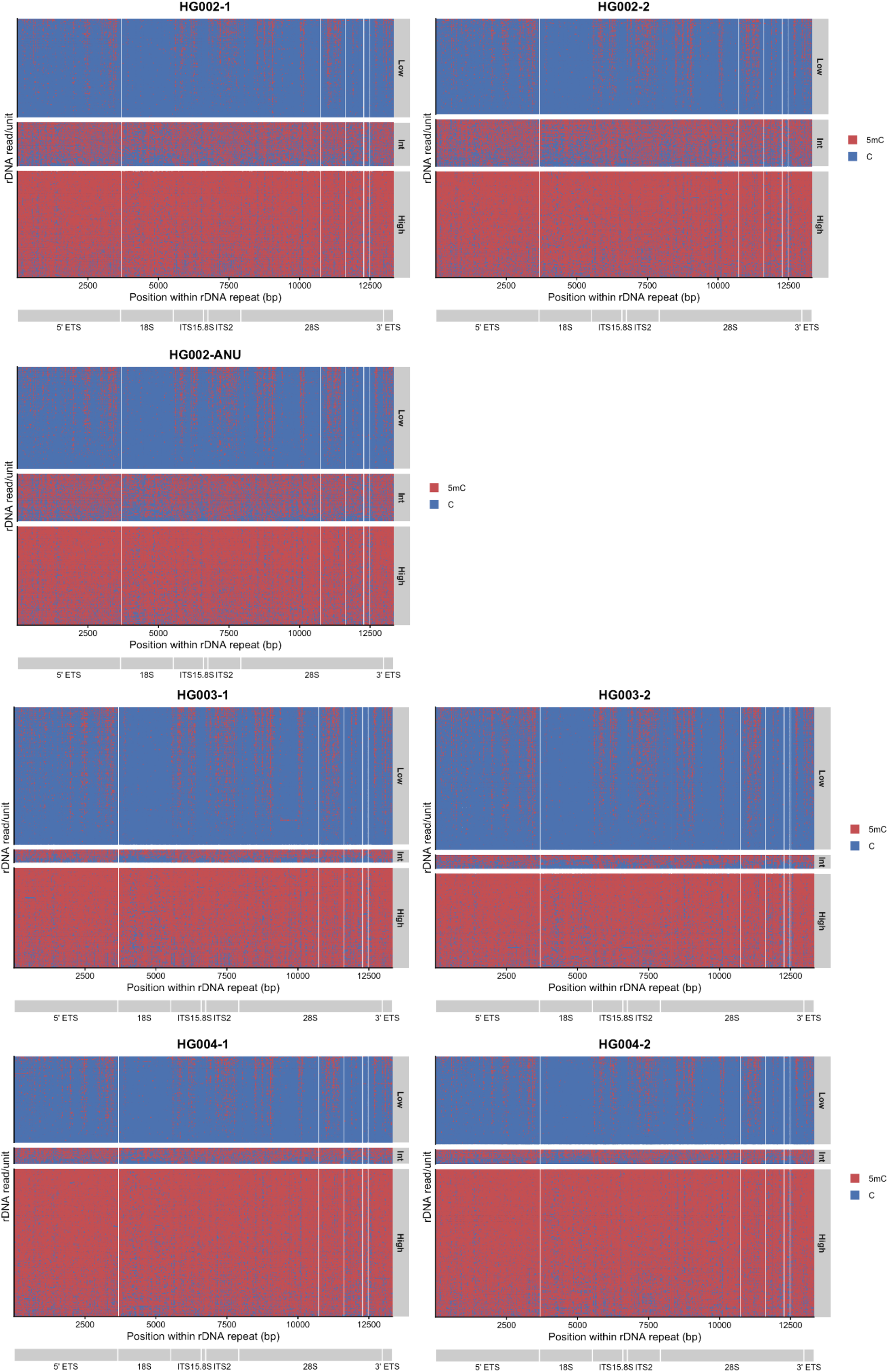
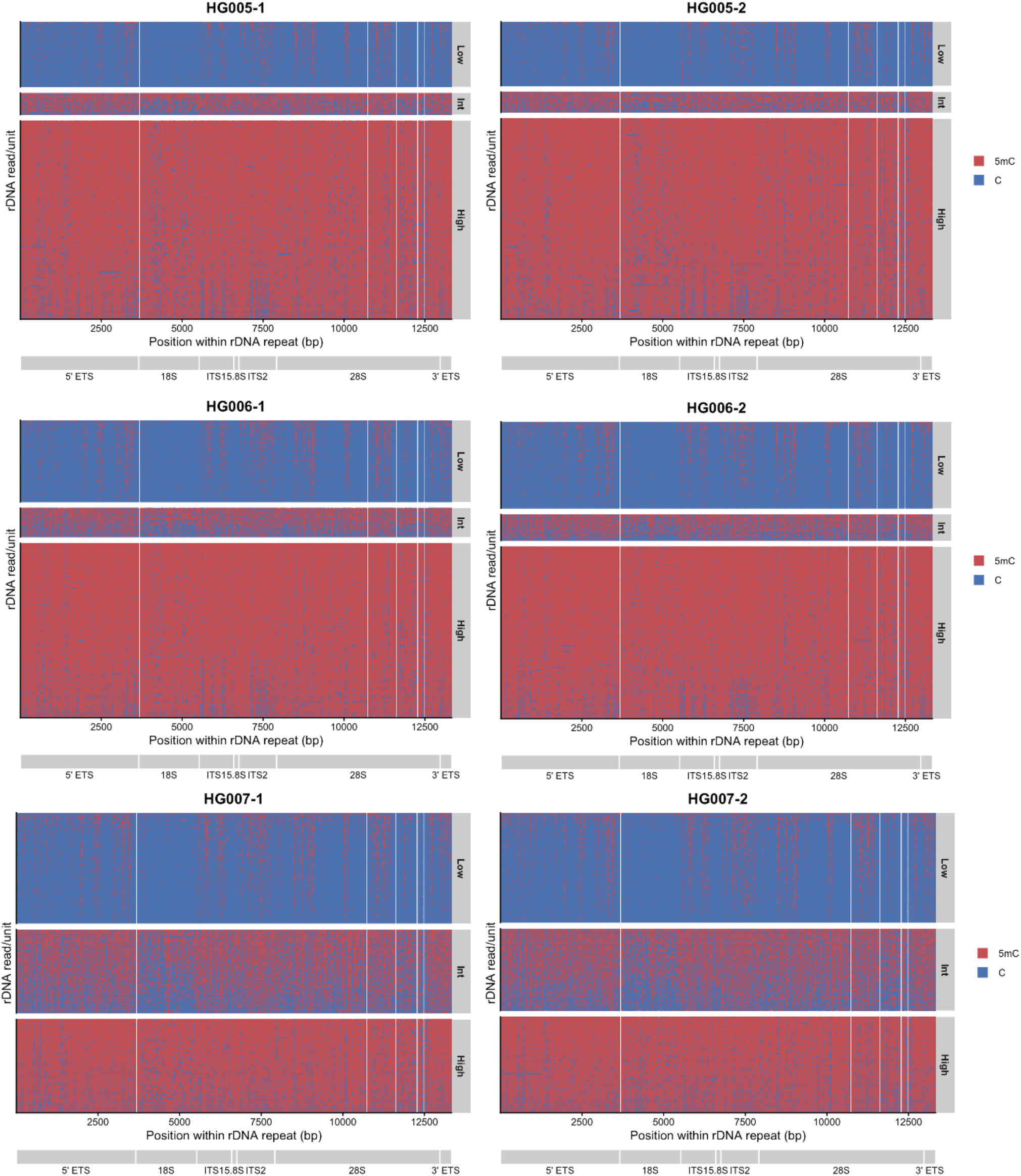
Spatial distribution of canonical C and 5mC calls across individual 45S rDNA repeats in the trio samples. Individual canonical C (blue) and 5mC (red) calls are shown according to their position within a single 45S rDNA repeat coordinate (1-13,332 bp). rDNA reads spanning multiple tandem repeat units were divided into individual read/repeat units, such that each row represents the portion of a sequencing read corresponding to one rDNA repeat unit rather than necessarily one independent sequencing read. Calls assigned to any of the five repeat units in the mapping reference were projected onto a common single-repeat coordinate. Read/repeat units were classified according to their overall 5mC proportion as low (<0.3), intermediate (0.3-0.7), or high (>0.7) methylation and grouped accordingly. Within each methylation class, read/repeat units are ordered by increasing 5mC proportion. The positions of the 5ʹ ETS, 18S, ITS1, 5.8S, ITS2, 28S and 3ʹ ETS regions are indicated below each plot. White regions represent positions without retained CpG calls for the corresponding read/repeat unit.

**Supplementary Figure S10.**
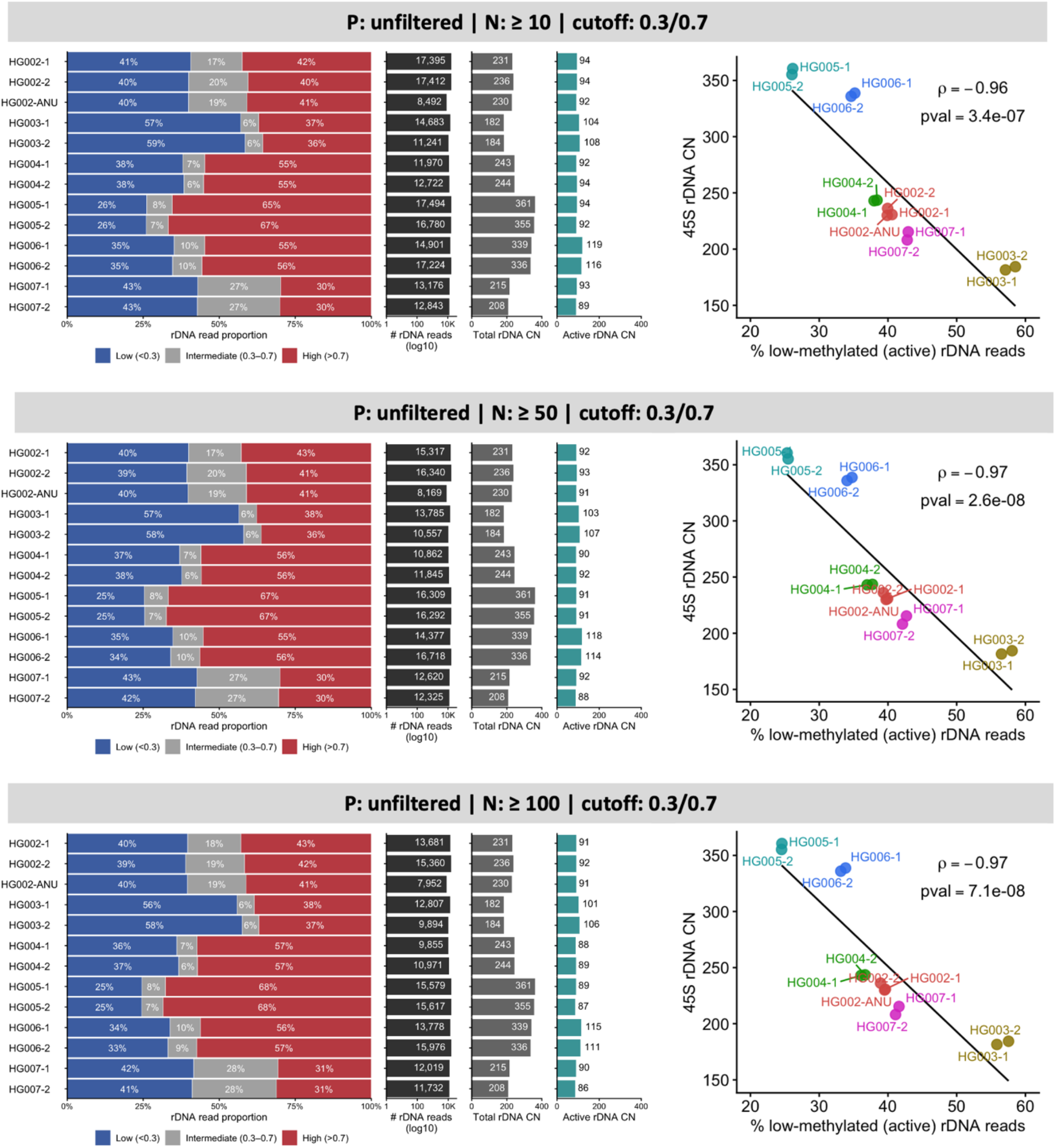
Per-read rDNA methylation profiles across trio samples using unfiltered 5mC calls and methylation classification cutoff of 0.3/0.7. The per-read methylation profiles on the left shows the proportion of reads classified as low (<0.3), intermediate (0.3-0.7), or high (>0.7) methylation based on the fraction of modified CpG sites per read, i.e. N_5mC_/(N_5mC_+N_C_). P denotes the per-base CpG calling-probability threshold and N denotes the minimum number of retained CpG sites required per rDNA read (N: ≥10, ≥50, ≥100). The accompanying panels show the total number of rDNA reads (log10 scale), estimated total rDNA CN, and estimated active rDNA CN calculated as the proportion of low-methylated reads. The condition (P: unfiltered | N: unfiltered | cutoff: 0.3/0.7) is shown in Fig. 4b. On the right, the regression plot shows the relationship between total rDNA CN (45S CN) and the proportion of low-methylated (active) rDNA reads across samples. Each point represents an individual run, as coloured by the sample. The black line shows the linear regression fit, with Spearman correlation (ρ) and corresponding p-value indicated.

**Supplementary Figure S11.**
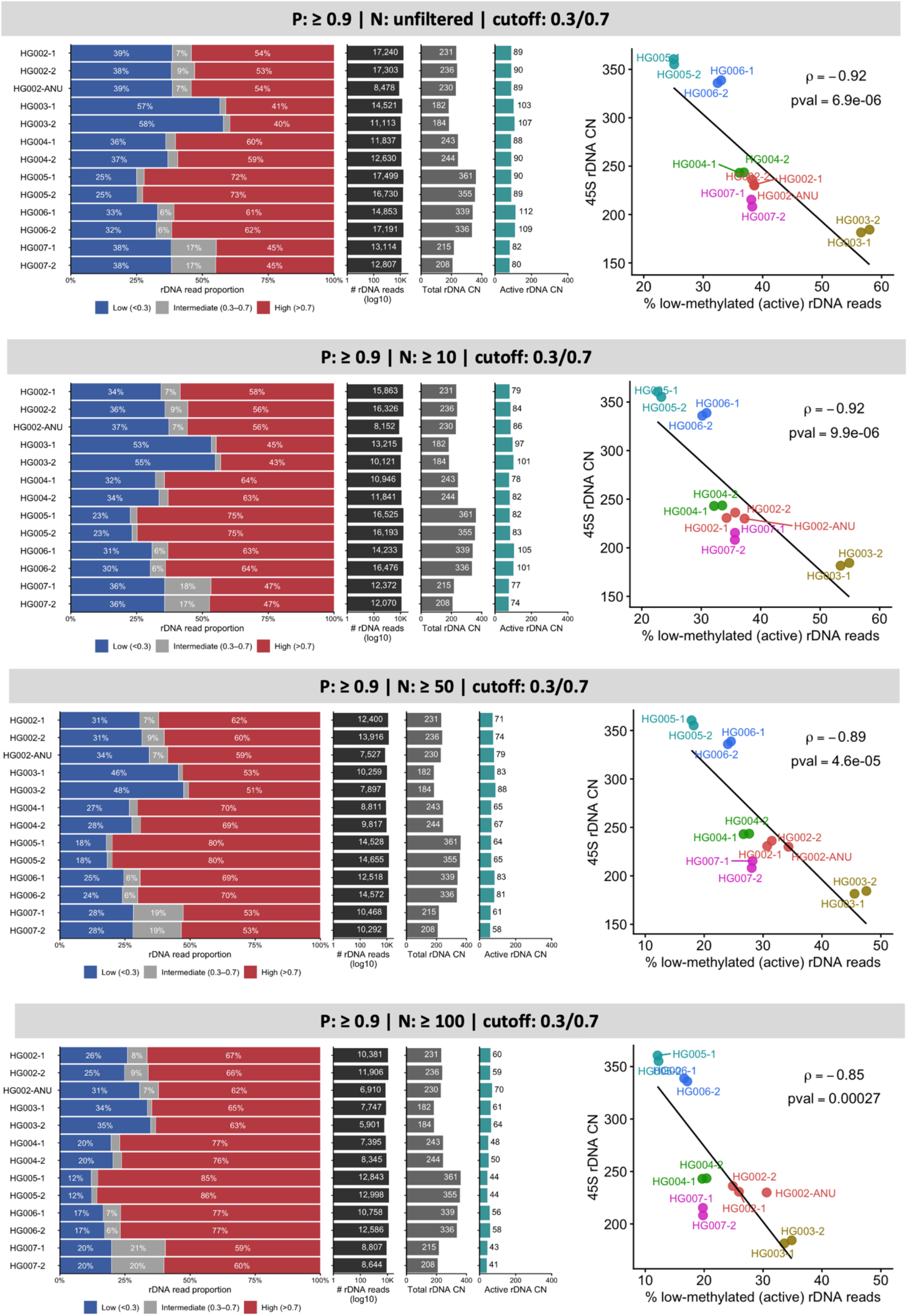
Per-read rDNA methylation profiles across trio samples using high-confidence 5mC calls (P ≥ 0.9) and methylation classification cutoff of 0.3/0.7. The per-read methylation profiles on the left shows the proportion of reads classified as low (<0.3), intermediate (0.3-0.7), or high (>0.7) methylation based on the fraction of modified CpG sites per read, i.e. N_5mC_/(N_5mC_+N_C_). P denotes the per-base CpG calling-probability threshold and N denotes the minimum number of retained CpG sites required per rDNA read (N: unfiltered, ≥10, ≥50, ≥100). The accompanying panels show the total number of rDNA reads (log10 scale), estimated total rDNA CN, and estimated active rDNA CN calculated as the proportion of low-methylated reads. The condition (P: unfiltered | N: unfiltered | cutoff: 0.3/0.7) is shown in Fig. 4b. On the right, the regression plot shows the relationship between total rDNA CN (45S CN) and the proportion of low-methylated (active) rDNA reads across samples. Each point represents an individual run, as coloured by the sample. The black line shows the linear regression fit, with Spearman correlation (ρ) and corresponding p-value indicated.

**Supplementary Figure S12.**
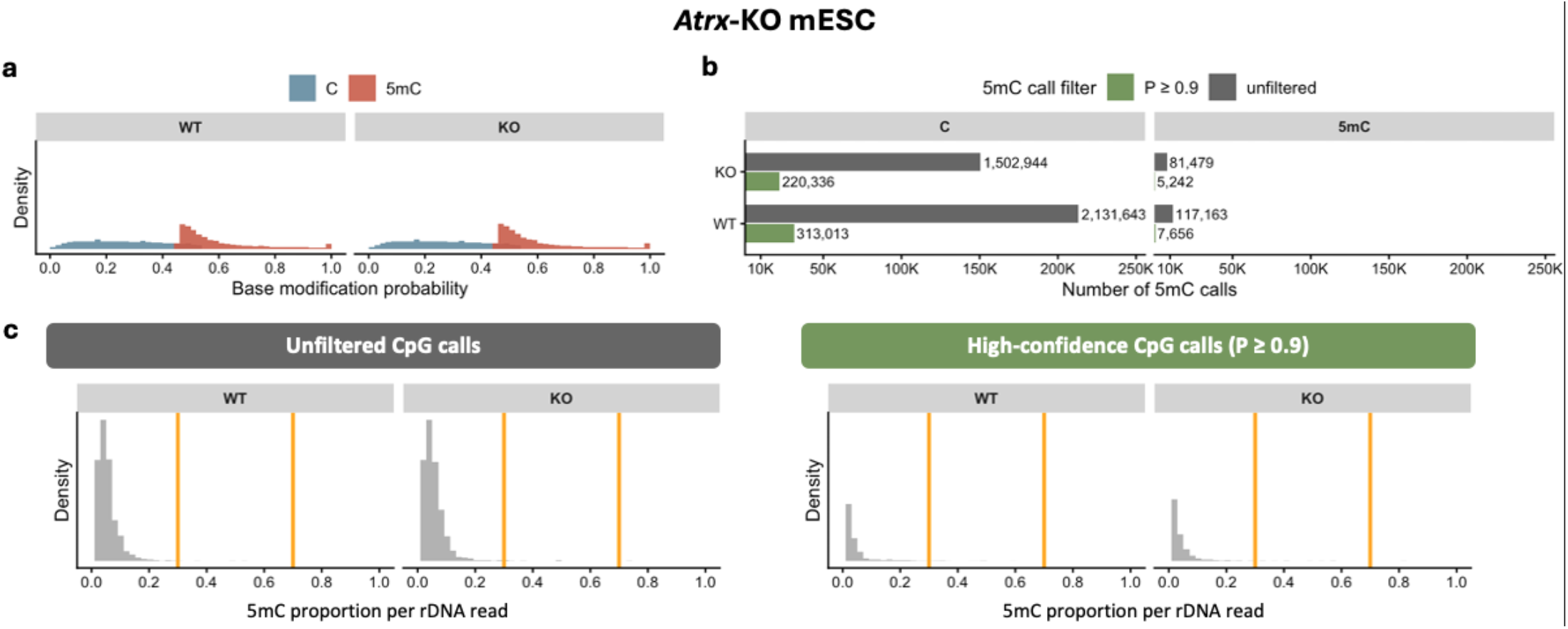
Base modification calling probabilities and read-level rDNA methylation distributions in wildtype (WT) and *Atrx*-knockout (KO) mouse embryonic stem cells (mESCs). **(a)** Distribution of base modification calling probabilities (P) for canonical C and 5mC calls across rDNA reads from WT and Atrx-KO mESCs. Canonical C probabilities are represented as the corresponding probability of modification (1 - P) for visualization on a common scale with 5mC calls **(b)** Number of canonical C (left) and 5mC (right) calls using unfiltered calls (dark grey) and after retaining high-confidence calls (P ≥ 0.9) (green) **(c)** Distribution of the proportion of 5mC calls per rDNA read using unfiltered CpG calls (left) or high-confidence CpG calls with P ≥ 0.9 (right), with no minimum number of CpG calls required per read. Orange lines indicate the 0.3 and 0.7 cutoffs used to classify reads as low-, intermediate-, and high-methylated, respectively.

**Supplementary Figure S13.**
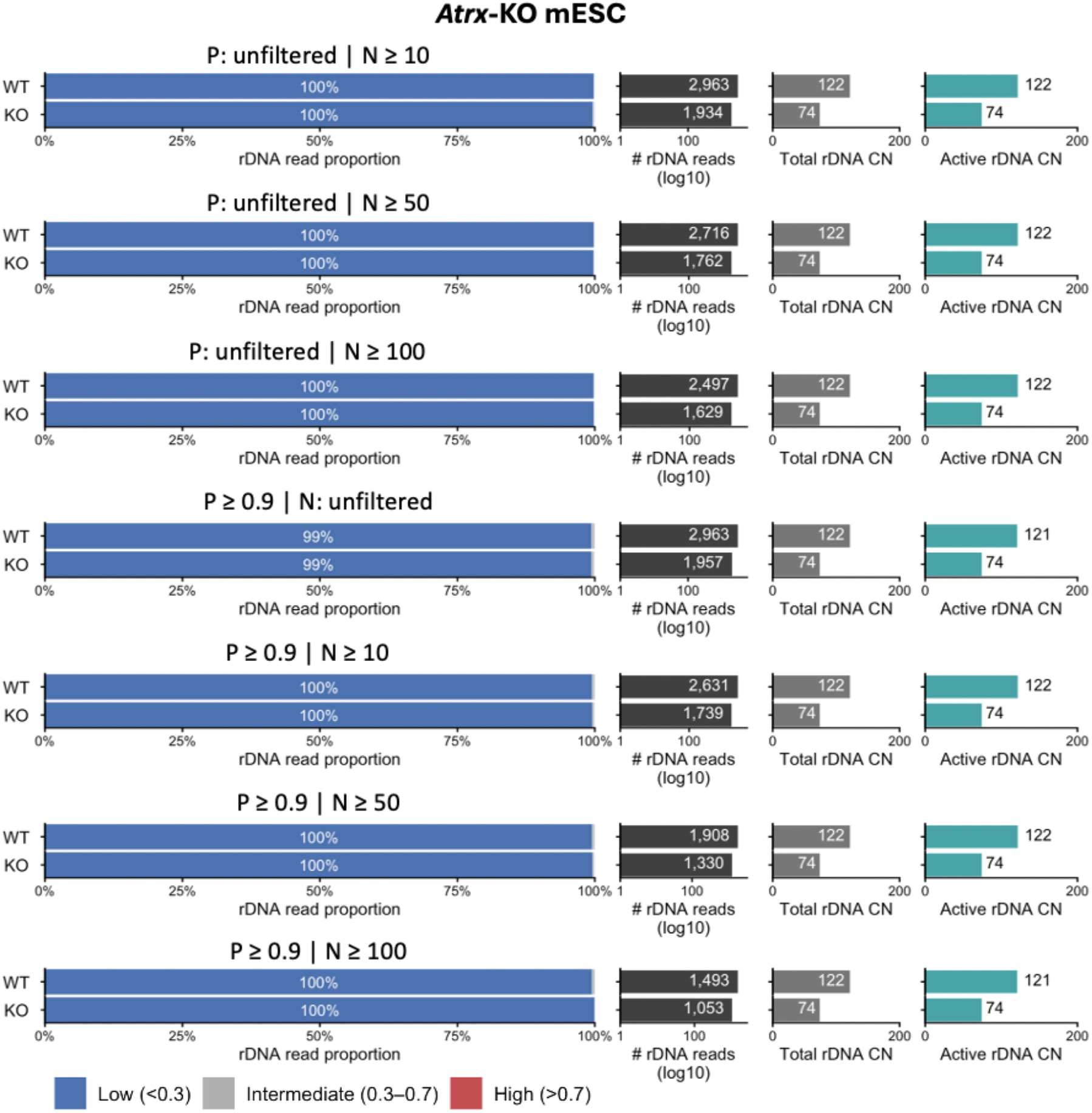
Robustness of rDNA methylation classification and putatively active rDNA copy number (CN) estimates to CpG filtering parameters in wildtype (WT) and *Atrx*-knockout (KO) mouse embryonic stem cells (mESCs). Per-read rDNA methylation profiles were evaluated using different combinations of 5mC calling probability filtering (P: unfiltered or P ≥ 0.9) and minimum number of CpG calls per rDNA read (N: unfiltered, ≥10, ≥50, or ≥100), with the 0.3/0.7 read-level methylation classification cutoffs. For each parameter combination, horizontal stacked bars show the proportions of rDNA reads classified as low (<0.3), intermediate (0.3–0.7), or high (>0.7) methylation. The number of retained rDNA reads in log10 scale, total rDNA CN, and estimated putatively active rDNA CN (total rDNA CN × fraction of low-methylated reads) are shown to the right.

**Supplementary Figure S14.**
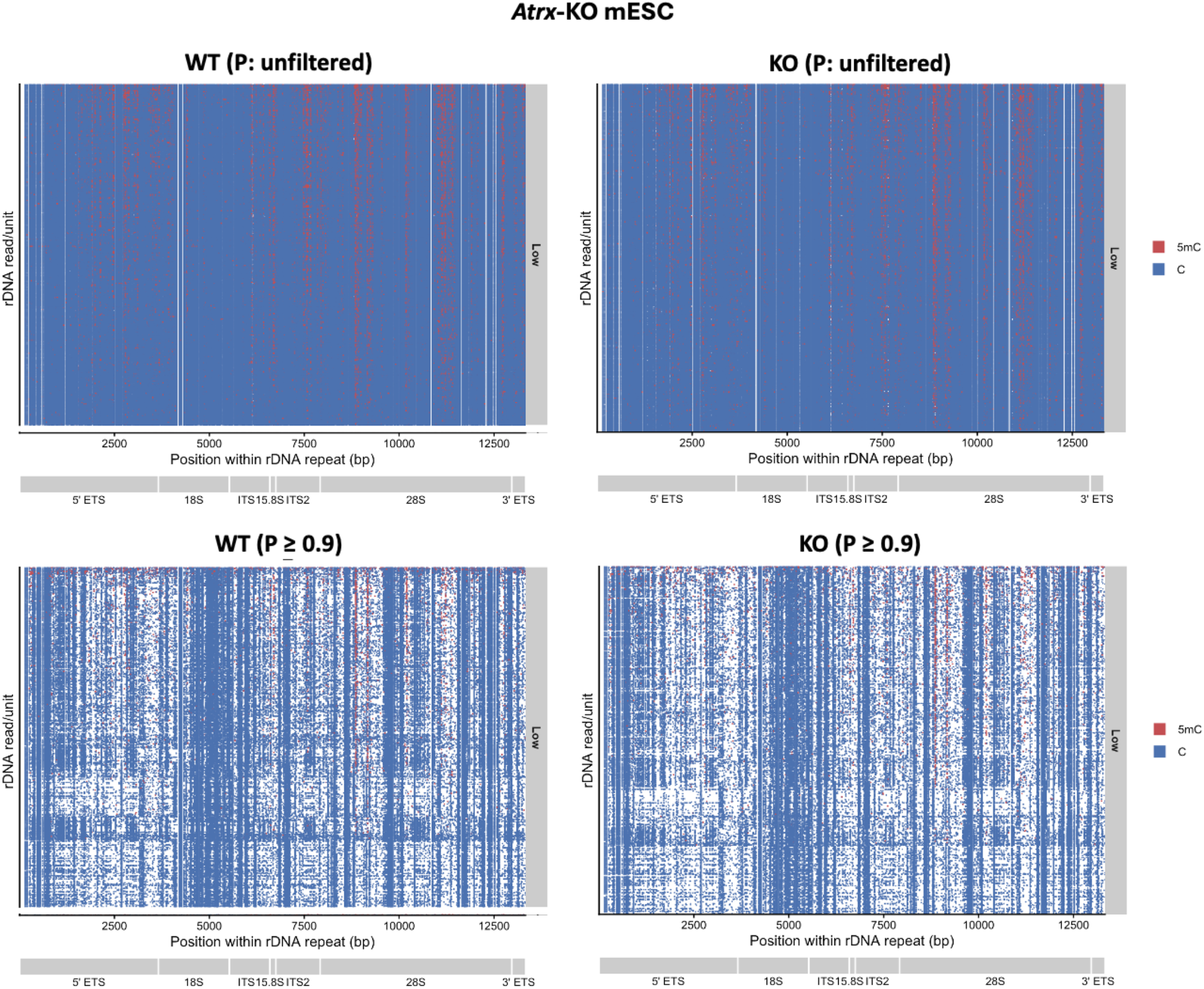
Spatial distribution of canonical C and 5mC calls across individual 45S rDNA repeats in the mouse embryonic stem cells (mESCs). Individual canonical C (blue) and 5mC (red) calls are shown according to their position within a single 45S rDNA repeat coordinate (1-13,332 bp). rDNA reads spanning multiple tandem repeat units were divided into individual read/repeat units, such that each row represents the portion of a sequencing read corresponding to one rDNA repeat unit rather than necessarily one independent sequencing read. Calls assigned to any of the five repeat units in the mapping reference were projected onto the one single repeat coordinate. Read/repeat units were classified according to their overall 5mC proportion as low (<0.3), intermediate (0.3-0.7), or high (>0.7) methylation and grouped accordingly, but all reads are classified as low methylation in this sample. The positions of the 5ʹ ETS, 18S, ITS1, 5.8S, ITS2, 28S and 3ʹ ETS regions are indicated below each plot. White regions represent positions without retained CpG calls for the corresponding read/repeat unit.

**Supplementary Figure S15.**
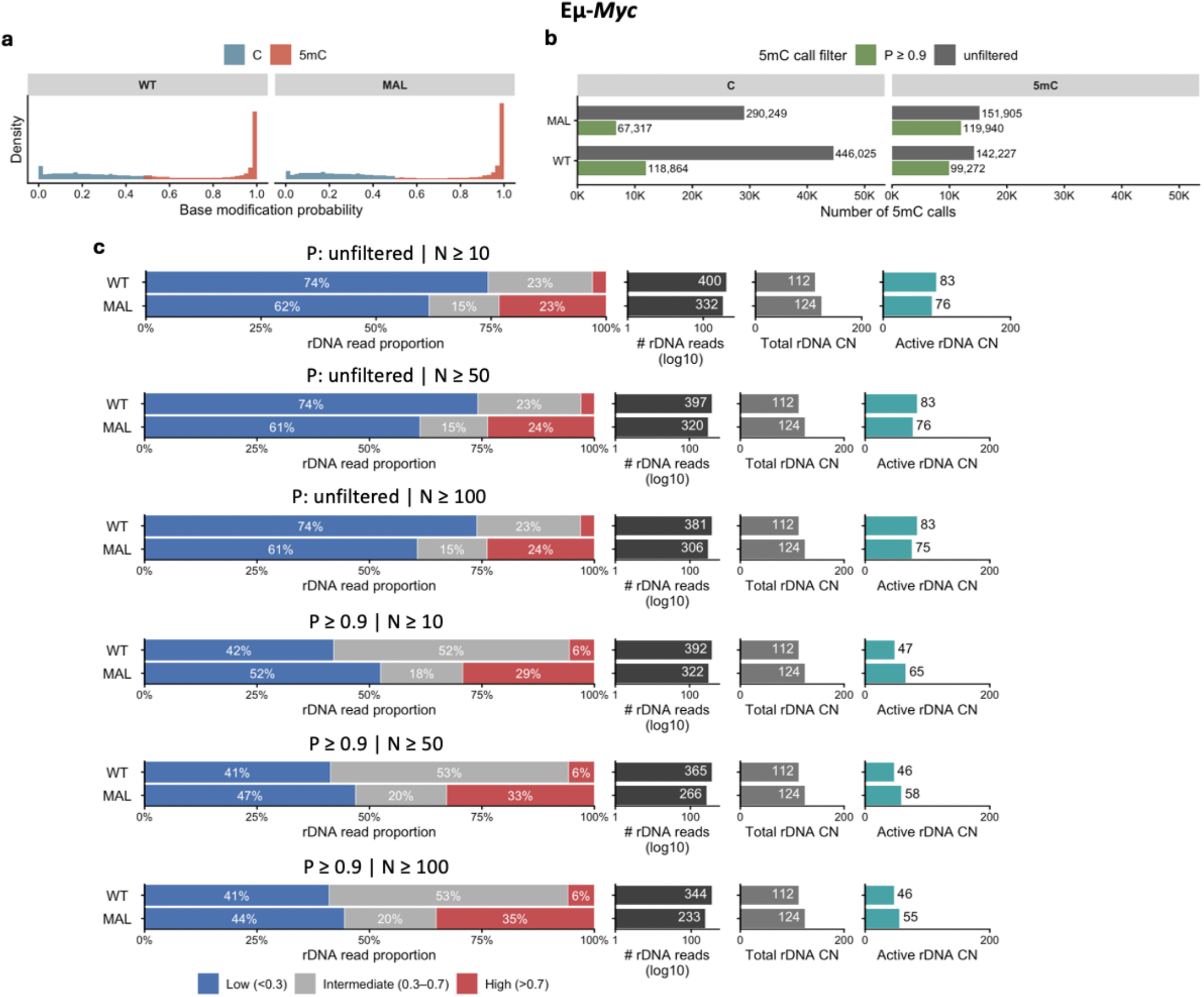
Base modification calling probabilities and read-level rDNA methylation distributions in wildtype (WT) and malignant (MAL) *Eμ-Myc* cells. **(a)** Distribution of base modification calling probabilities (P) for canonical C and 5mC calls across rDNA reads from WT and MAL Eμ-*Myc* cells. Canonical C probabilities are represented as the corresponding probability of modification (1 - P) for visualization on a common scale with 5mC calls. **(b)** Number of canonical C (left) and 5mC (right) calls before call-probability filtering (i.e. unfiltered) (dark grey) and after retaining high-confidence calls (P ≥ 0.9) (green). **(c)** Per-read rDNA methylation profiles were evaluated using different combinations of 5mC calling probability filtering (P: unfiltered or P ≥ 0.9) and minimum numbers of CpG calls per rDNA read (N: unfiltered, ≥10, ≥50, or ≥100), with the 0.3/0.7 read-level methylation classification cutoffs. For each parameter combination, horizontal stacked bars show the proportions of rDNA reads classified as low (<0.3), intermediate (0.3–0.7), or high (>0.7) methylation. The number of retained rDNA reads in log10 scale, total rDNA CN, and estimated putatively active rDNA CN (total rDNA CN × fraction of low-methylated reads) are shown to the right.

**Supplementary Figure S16.**
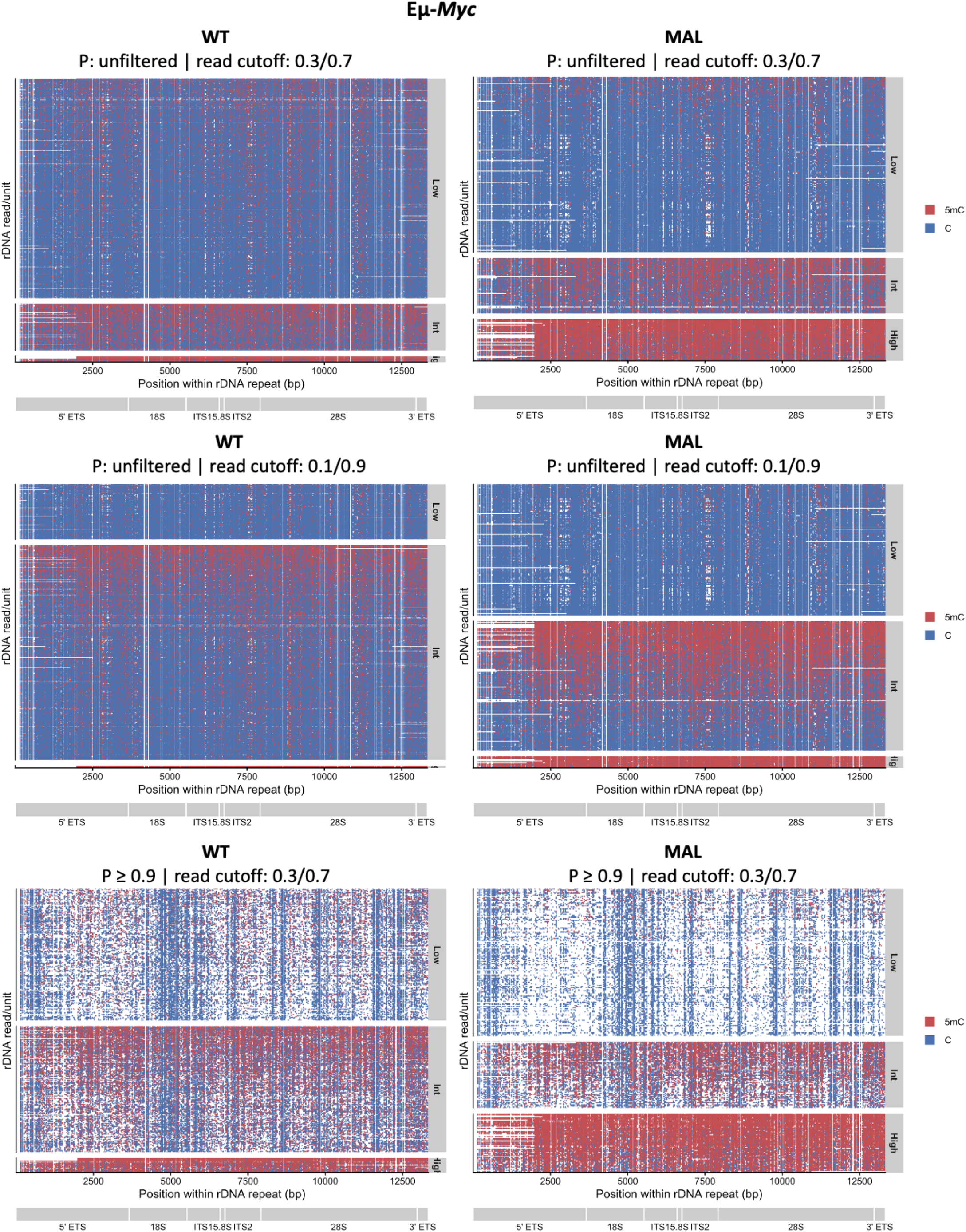
Spatial distribution of canonical C and 5mC calls across individual 45S rDNA repeats in wildtype (WT) and malignant (MAL) *Eμ-Myc* cells. Individual canonical C (blue) and 5mC (red) calls are shown according to their position within a single 45S rDNA repeat coordinate (1-13,332 bp). rDNA reads spanning multiple tandem repeat units were divided into individual read/repeat units, such that each row represents the portion of a sequencing read corresponding to one rDNA repeat unit rather than necessarily one independent sequencing read. Calls assigned to any of the five repeat units in the mapping reference were projected onto a common single-repeat coordinate. Read/repeat units were generated using either unfiltered CpG calls or high-confidence CpG calls (P ≥ 0.9), and were classified according to their overall 5mC proportion using either the 0.3/0.7 or 0.1/0.9 read-level methylation cutoffs, as indicated above each panel. The positions of the 5ʹ ETS, 18S, ITS1, 5.8S, ITS2, 28S and 3ʹ ETS regions are indicated below each plot. White regions represent positions without retained CpG calls for the corresponding read/repeat unit.

**Supplementary Figure S17.**
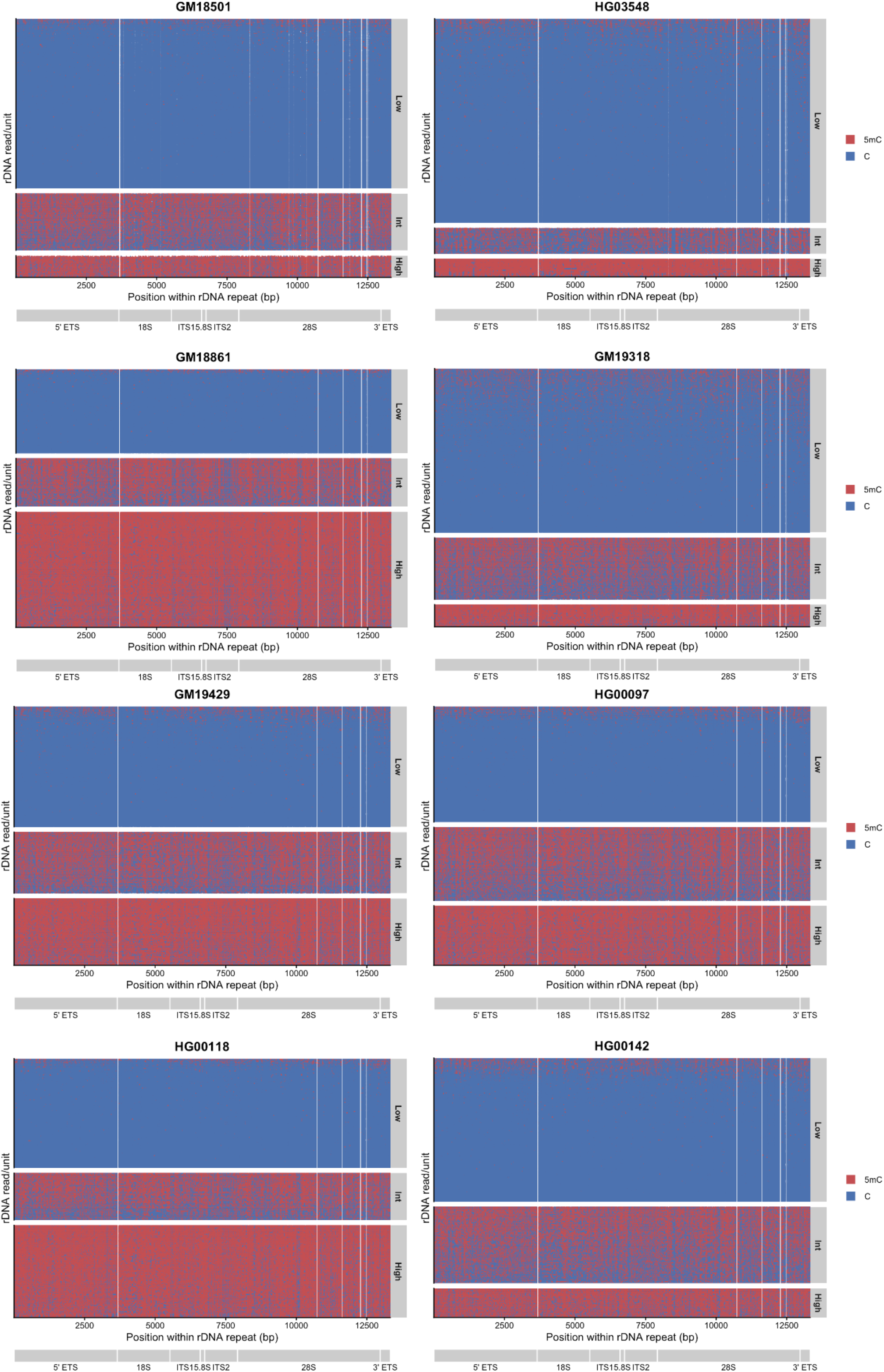
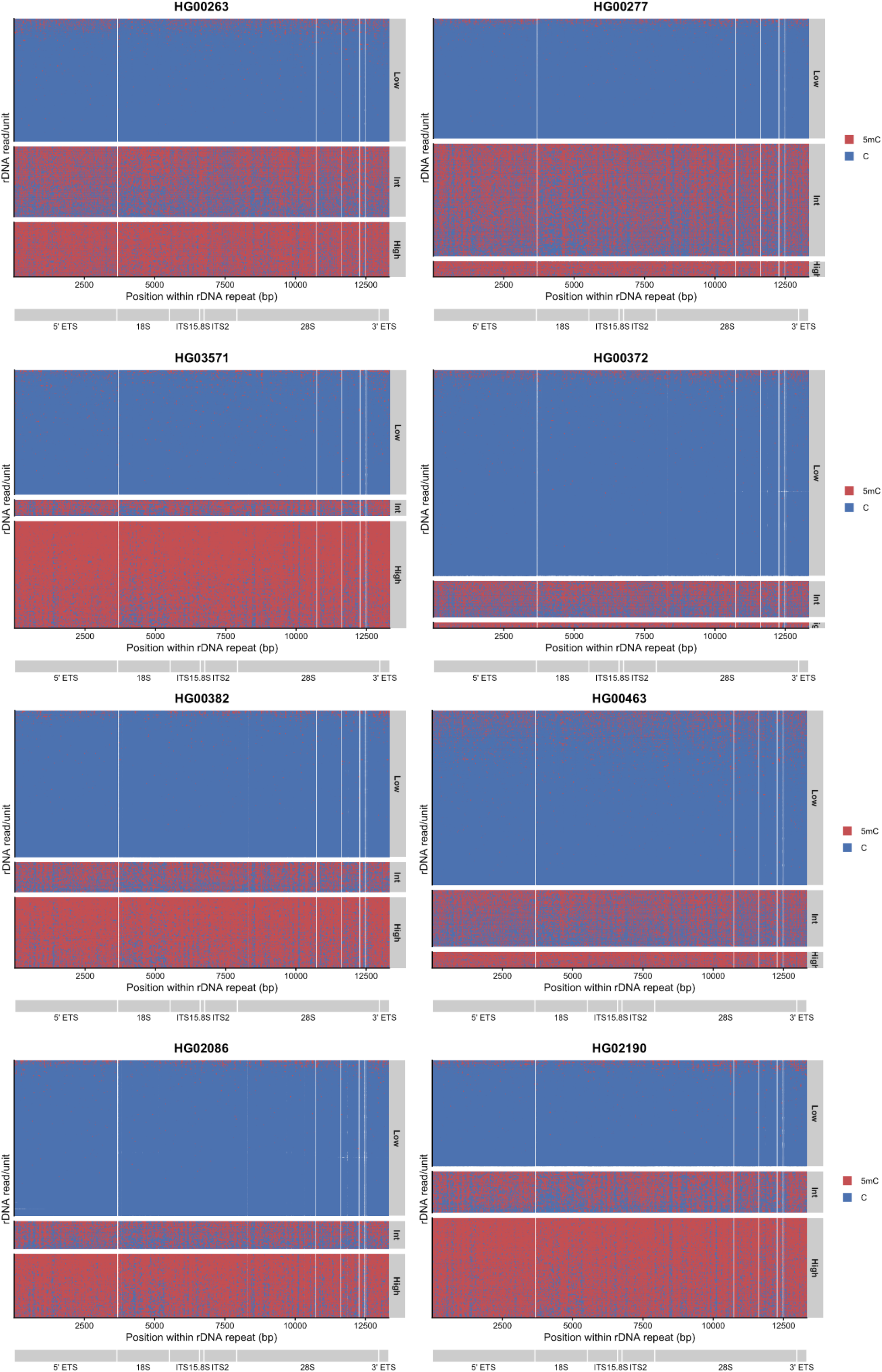
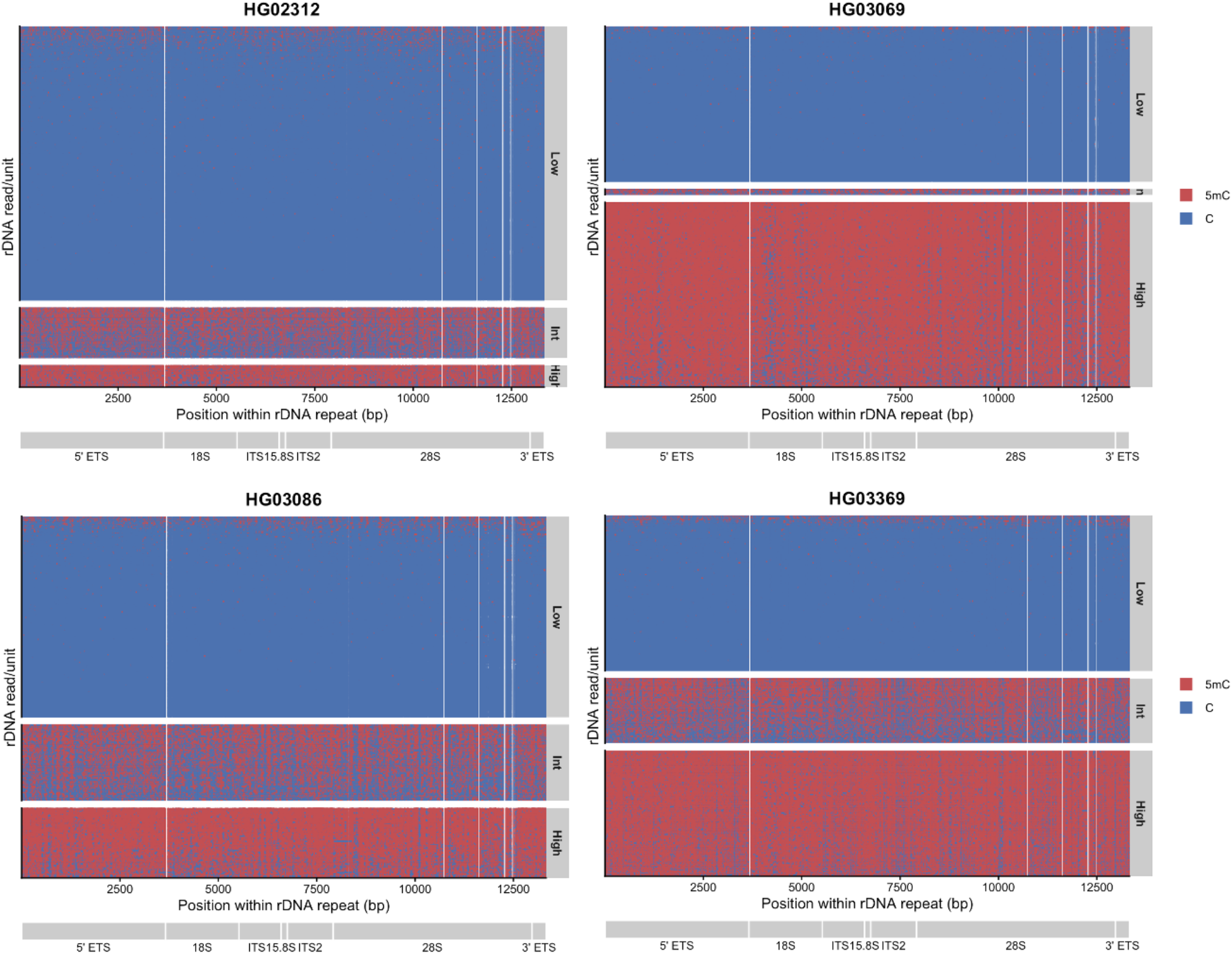
Spatial distribution of canonical C and 5mC calls across individual 45S rDNA repeats in 1KGP samples. Individual canonical C (blue) and 5mC (red) calls are shown according to their positions within a single 45S rDNA repeat coordinate (1-13,332 bp). rDNA reads spanning multiple tandem repeat units were divided into individual read/repeat units, such that each row represents the portion of a sequencing read corresponding to one rDNA repeat unit rather than necessarily one independent sequencing read. Calls assigned to any of the five repeat units in the mapping reference were projected onto a common single-repeat coordinate. Read/repeat units were classified according to their overall 5mC proportion as low (<0.3), intermediate (0.3-0.7), or high (>0.7) methylation and grouped accordingly. The positions of the 45S regions are indicated below each plot. White regions represent positions without CpG calls for the corresponding read/repeat unit.

**Supplementary Figure S18.**
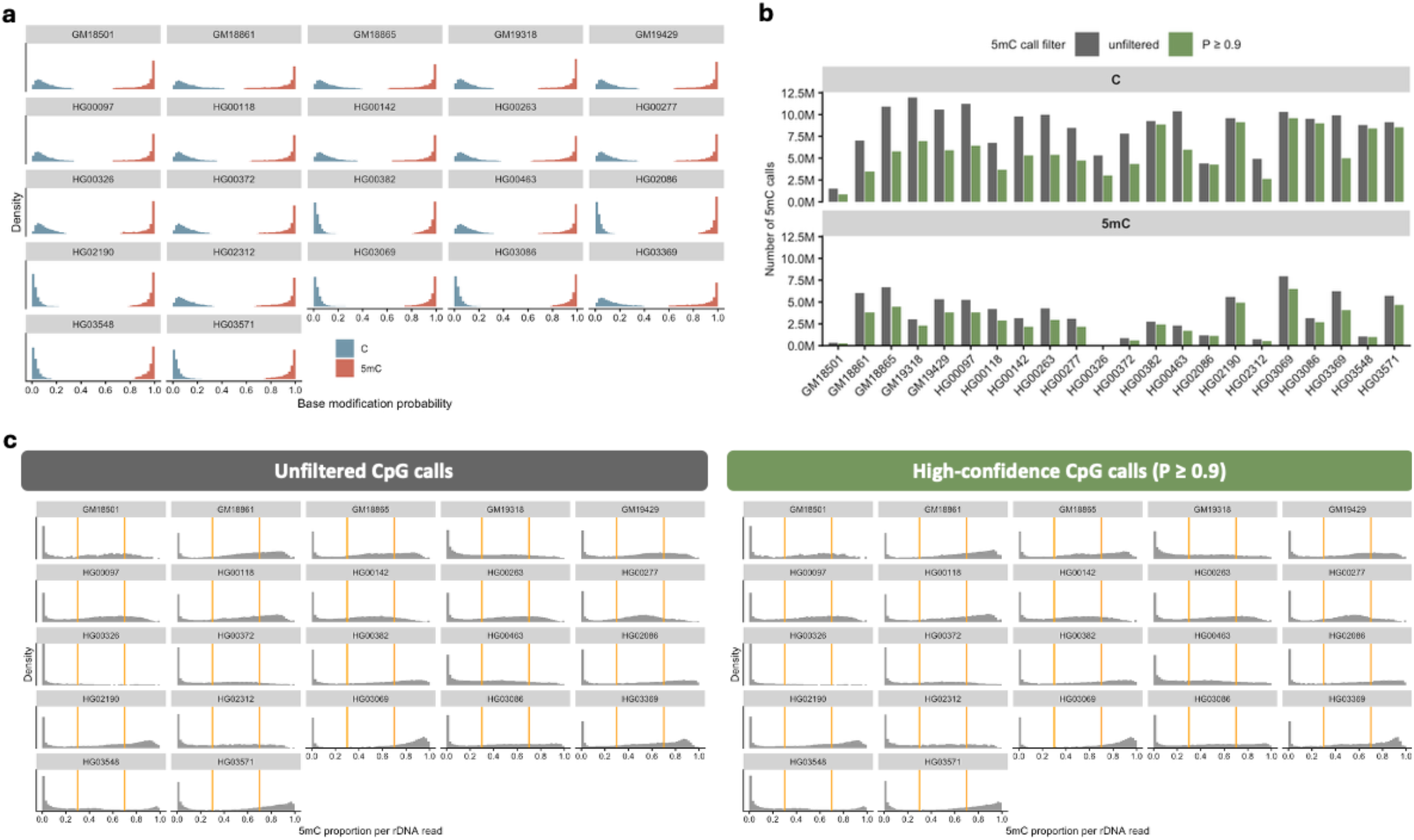
Base modification calling probabilities and read-level rDNA methylation distributions across 1KGP samples. **(a)** Distribution of base modification calling probabilities (P) for canonical C and 5mC calls across rDNA reads across 22 1KGP samples. Canonical C probabilities are represented as the corresponding probability of modification (1 - P) for visualization on a common scale with 5mC calls. **(b)** Number of canonical C and 5mC calls before call-probability filtering (unfiltered) and after retaining high-confidence calls with P ≥ 0.9. **(c)** Distribution of the proportion of 5mC calls per rDNA read using unfiltered CpG calls (left) or high-confidence CpG calls with P ≥ 0.9 (right), with no minimum number of CpG calls required per read. Orange lines indicate the 0.3 and 0.7 cutoffs used to classify reads as low-, intermediate-, and high-methylated, respectively.

**Supplementary Figure S19.**
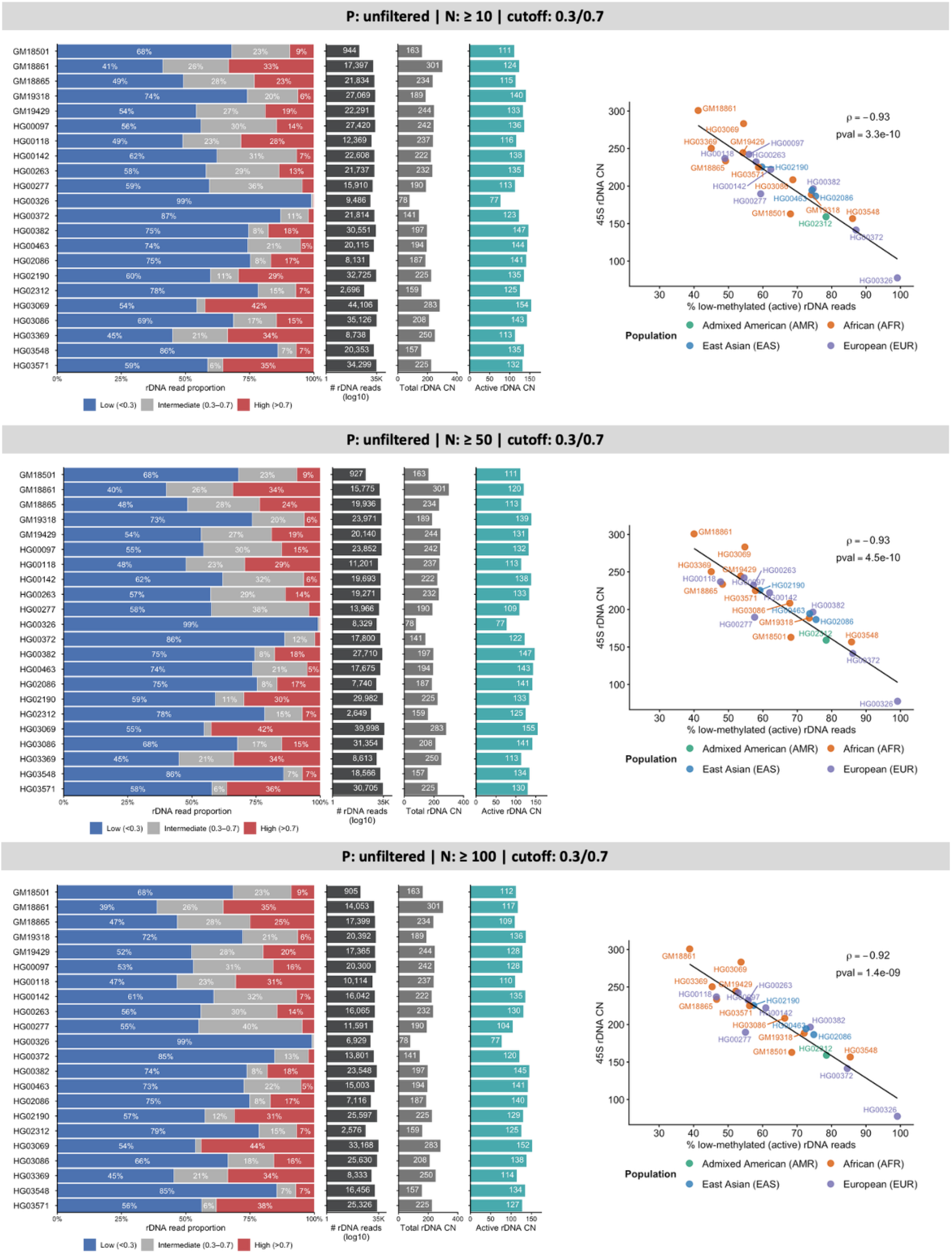
Robustness of rDNA methylation classification and putatively active rDNA CN estimates to minimum CpG per read filtering across 1KGP samples using unfiltered CpG calls. Per-read rDNA methylation profiles were evaluated using unfiltered CpG calls and minimum numbers of CpG calls per rDNA read (*N*) of ≥10, ≥50, or ≥100, with the 0.3/0.7 read-level methylation classification cutoffs. For each condition, horizontal stacked bars show the proportions of rDNA reads classified as low (<0.3), intermediate (0.3-0.7), or high (>0.7) methylation. The number of retained rDNA reads (log10 scale), total rDNA CN, and estimated putatively active rDNA CN (total rDNA CN × fraction of low-methylated reads) are shown in the adjacent panels. Regression plots show the relationship between total rDNA CN and the proportion of low-methylated rDNA reads under each filtering condition. Points are coloured by population affiliation, and the black line shows the linear regression fit. Spearman’s correlation coefficient (ρ) and corresponding *P* value are indicated. The corresponding analysis with no minimum CpG per read requirement is shown in Fig. 6a & b.

**Supplementary Figure S20.**
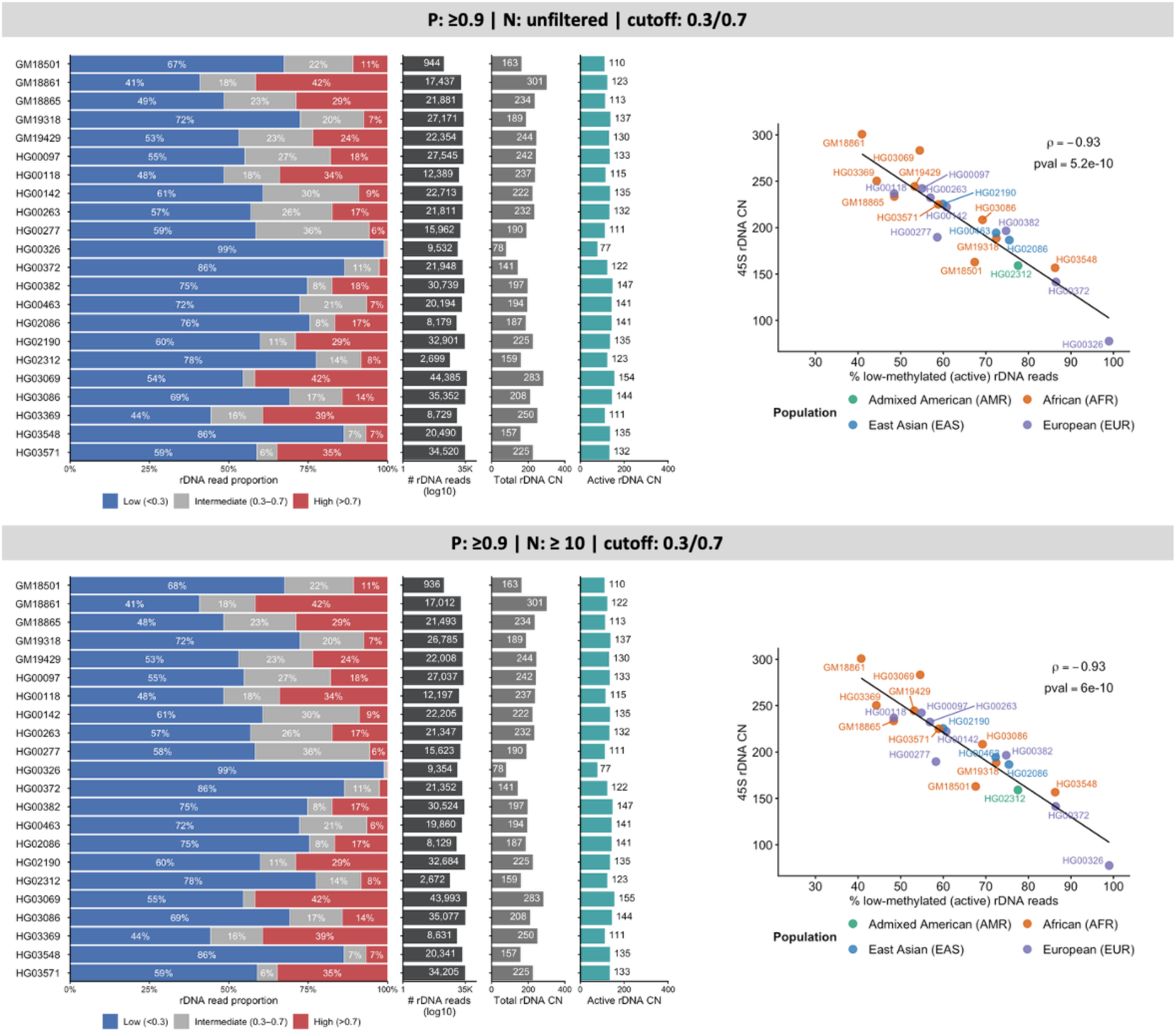
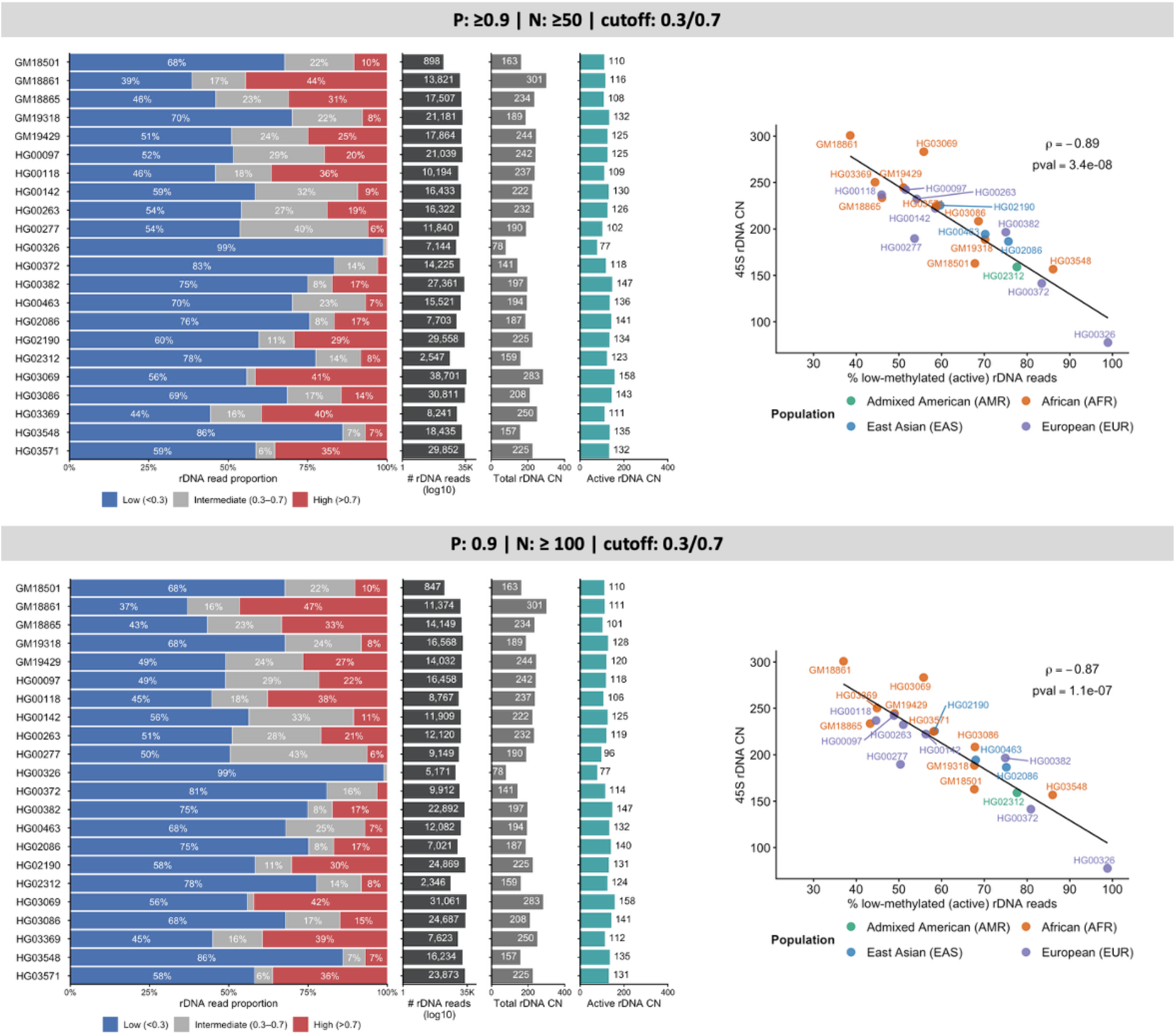
Robustness of rDNA methylation classification and putatively active rDNA CN estimates to minimum CpG per read filtering across 1KGP samples using high-confidence CpG calls. Per-read rDNA methylation profiles were evaluated using high-confidence CpG calls (*P* ≥ 0.9) with either no minimum number of CpG calls per rDNA read or minimum requirements (*N*) of ≥10, ≥50, or ≥100, using the 0.3/0.7 read-level methylation classification thresholds. For each condition, horizontal stacked bars show the proportions of rDNA reads classified as low (<0.3), intermediate (0.3-0.7), or high (>0.7) methylation. The number of retained rDNA reads (log10 scale), total rDNA CN, and estimated putatively active rDNA CN (total rDNA CN × fraction of low-methylated reads) are shown in the adjacent panels. Regression plots show the relationship between total rDNA CN and the proportion of low-methylated rDNA reads under each filtering condition. Points are coloured by population affiliation, and the black line shows the linear regression fit. Spearman’s correlation coefficient (ρ) and corresponding *P* value are indicated.

## References

1. Betto, R. M., Diamante, L., Perrera, V., Audano, M., Rapelli, S., Lauria, A., . . . Martello, G. (2021). Metabolic control of DNA methylation in naive pluripotent cells. Nature genetics, 53(2), 215–229. doi:10.1038/s41588-020-00770-2

2. Britton-Davidian, J., Cazaux, B., & Catalan, J. (2012). Chromosomal dynamics of nucleolar organizer regions (NORs) in the house mouse: micro-evolutionary insights. Heredity (Edinb), 108(1), 68–74. doi:10.1038/hdy.2011.105

3. Bywater, M. J., Poortinga, G., Sanij, E., Hein, N., Peck, A., Cullinane, C., . . . Anderes, K. (2012). Inhibition of RNA polymerase I as a therapeutic strategy to promote cancer-specific activation of p53. Cancer cell, 22(1), 51–65.

4. Diesch, J., Bywater, M. J., Sanij, E., Cameron, D. P., Schierding, W., Brajanovski, N., . . . Poortinga, G. (2019). Changes in long-range rDNA-genomic interactions associate with altered RNA polymerase II gene programs during malignant transformation. Commun Biol, 2, 39. doi:10.1038/s42003-019-0284-y

5. Fan, W., Eklund, E., Sherman, R. M., Liu, H., Pitts, S., Ford, B., . . . Laiho, M. (2022). Widespread genetic heterogeneity of human ribosomal RNA genes. RNA, 28(4), 478–492.

6. Gagnon-Kugler, T., Langlois, F., Stefanovsky, V., Lessard, F., & Moss, T. (2009). Loss of human ribosomal gene CpG methylation enhances cryptic RNA polymerase II transcription and disrupts ribosomal RNA processing. Molecular cell, 35(4), 414–425.

7. Gál, Z., Boukoura, S., Oxe, K. C., Badawi, S., Nieto, B., Korsholm, L. M., . . . Dahl, C. (2024). Hyper-recombination in ribosomal DNA is driven by long-range resection-independent RAD51 accumulation. Nature Communications, 15(1), 7797.

8. Gustafson, J. A., Gibson, S. B., Damaraju, N., Zalusky, M. P. G., Hoekzema, K., Twesigomwe, D., . . . Miller, D. E. (2024). High-coverage nanopore sequencing of samples from the 1000 Genomes Project to build a comprehensive catalog of human genetic variation. Genome Res, 34(11), 2061–2073. doi:10.1101/gr.279273.124

9. Hall, A. N., Turner, T. N., & Queitsch, C. (2021). Thousands of high-quality sequencing samples fail to show meaningful correlation between 5S and 45S ribosomal DNA arrays in humans. Scientific reports, 11(1), 449. doi:10.1038/s41598-020-80049-y

10. Hallgren, J., Pietrzak, M., Rempala, G., Nelson, P. T., & Hetman, M. (2014). Neurodegeneration-associated instability of ribosomal DNA. Biochimica et Biophysica Acta (BBA)-Molecular Basis of Disease, 1842(6), 860–868.

11. Hein, N., Hannan, K. M., George, A. J., Sanij, E., & Hannan, R. D. (2013). The nucleolus: an emerging target for cancer therapy. Trends in molecular medicine, 19(11), 643–654.

12. Henderson, A., Warburton, D., & Atwood, K. (1972). Location of ribosomal DNA in the human chromosome complement. Proceedings of the National Academy of Sciences, 69(11), 3394–3398.

13. Hori, Y., Engel, C., & Kobayashi, T. (2023). Regulation of ribosomal RNA gene copy number, transcription and nucleolus organization in eukaryotes. Nature Reviews Molecular Cell Biology, 24(6), 414–429.

14. Hori, Y., Shimamoto, A., & Kobayashi, T. (2021). The human ribosomal DNA array is composed of highly homogenized tandem clusters. Genome Research, 31(11), 1971–1982.

15. McStay, B. (2016). Nucleolar organizer regions: genomic ‘dark matter’requiring illumination. Genes & development, 30(14), 1598–1610.

16. McStay, B., & Grummt, I. (2008). The epigenetics of rRNA genes: from molecular to chromosome biology. Annual review of cell and developmental biology, 24, 131–157.

17. Moss, T., Langlois, F., Gagnon-Kugler, T., & Stefanovsky, V. (2007). A housekeeper with power of attorney: the rRNA genes in ribosome biogenesis. Cellular and Molecular Life Sciences, 64(1), 29–49. doi:10.1007/s00018-006-6278-1

18. Nurk, S., Koren, S., Rhie, A., Rautiainen, M., Bzikadze, A. V., Mikheenko, A., . . . Gershman, A. (2022). The complete sequence of a human genome. Science, 376(6588), 44–53.

19. Panov, K. I., Hannan, K., Hannan, R. D., & Hein, N. (2021). The Ribosomal Gene Loci-The Power behind the Throne. Genes (Basel), 12(5). doi:10.3390/genes12050763

20. Parks, M. M., Kurylo, C. M., Dass, R. A., Bojmar, L., Lyden, D., Vincent, C. T., & Blanchard, S. C. (2018). Variant ribosomal RNA alleles are conserved and exhibit tissue-specific expression. Science Advances, 4(2), eaao0665.

21. Rodriguez-Algarra, F., Evans, D. M., & Rakyan, V. K. (2024). Ribosomal DNA copy number variation associates with hematological profiles and renal function in the UK Biobank. Cell Genomics, 4(6).

22. Rodriguez-Algarra, F., Seaborne, R. A. E., Danson, A. F., Yildizoglu, S., Yoshikawa, H., Law, P. P., . . . Rakyan, V. K. (2022). Genetic variation at mouse and human ribosomal DNA influences associated epigenetic states. Genome Biol, 23(1), 54. doi:10.1186/s13059-022-02617-x

23. Rodriguez-Algarra, F., Whittaker, E., del Castillo del Rio, S., & Rakyan, V. K. (2025). Assessing human ribosomal DNA variation and its association with phenotypic outcomes. Bioessays, 47(4), e202400232.

24. Schlesinger, S., Selig, S., Bergman, Y., & Cedar, H. (2009). Allelic inactivation of rDNA loci. Genes & development, 23(20), 2437–2447.

25. Sharma, D., Denmat, S. H.-L., Matzke, N. J., Hannan, K., Hannan, R. D., O’Sullivan, J. M., & Ganley, A. R. D. (2022). A new method for determining ribosomal DNA copy number shows differences between Saccharomyces cerevisiae populations. Genomics, 114(4), 110430. 10.1016/j.ygeno.2022.110430

26. Udugama, M., Sanij, E., Voon, H. P., Son, J., Hii, L., Henson, J. D., . . . Pearson, R. B. (2018). Ribosomal DNA copy loss and repeat instability in ATRX-mutated cancers. Proceedings of the National Academy of Sciences, 115(18), 4737–4742.

27. Valori, V., Tus, K., Laukaitis, C., Harris, D. T., LeBeau, L., & Maggert, K. A. (2020). Human rDNA copy number is unstable in metastatic breast cancers. Epigenetics, 15(1-2), 85–106. doi:10.1080/15592294.2019.1649930

28. Wick, R. R. (2019). Badread: simulation of error-prone long reads. Journal of Open Source Software, 4(36), 1316.

